# Sequential structure probing of cotranscriptional RNA folding intermediates

**DOI:** 10.1101/2024.10.14.618260

**Authors:** Courtney E. Szyjka, Skyler L. Kelly, Eric J. Strobel

**Affiliations:** Department of Biological Sciences, The University at Buffalo, Buffalo, NY 14260, USA

## Abstract

Cotranscriptional RNA folding pathways typically involve the sequential formation of folding intermediates. Existing methods for cotranscriptional RNA structure probing map the structure of nascent RNA in the context of a terminally arrested transcription elongation complex. Consequently, the rearrangement of RNA structures as nucleotides are added to the transcript can be inferred but is not assessed directly. To address this limitation, we have developed linked-<u>m</u>ultipoint <u>T</u>ranscription <u>E</u>longation <u>C</u>omplex RNA structure <u>prob</u>ing (TECprobe-LM), which assesses the cotranscriptional rearrangement of RNA structures by sequentially positioning *E. coli* RNAP at two or more points within a DNA template so that nascent RNA can be chemically probed. We validated TECprobe-LM by measuring known folding events that occur within the *E. coli* signal recognition particle RNA, *Clostridium beijerinckii pfl* ZTP riboswitch, and *Bacillus cereus crcB* fluoride riboswitch folding pathways. Our findings establish TECprobe-LM as a strategy for detecting cotranscriptional RNA folding events directly using chemical probing.

## Introduction

RNA begins to fold cotranscriptionally as it is synthesized by an RNA polymerase (RNAP)^1–3^. Because base pair formation occurs ∼3 orders of magnitude faster than nucleotide addition, nascent RNA structures can begin to fold once RNA has emerged from the RNA polymerase footprint^4,5^. Consequently, cotranscriptional RNA folding pathways typically comprise a sequence of folding intermediates, some of which may be transient structures that do not persist within the native structure of the full-length RNA^6–11^.

In the past decade, biochemical and biophysical methods for measuring cotranscriptional RNA structure and folding have begun to enable the detection of RNA folding intermediates and the reconstruction of RNA folding pathways from data^9–20^. Among the experimental approaches that have been developed, cotranscriptional RNA structure probing applies high-throughput RNA chemical probing to measure the structure of nascent RNA. Existing cotranscriptional RNA structure probing methods capture RNA folding intermediates by systematically arresting RNAP at each position of a DNA template so that cotranscriptionally folded RNA is displayed from a static transcription elongation complex (TEC) and can be chemically probed^9,10,13,20^. This strategy can be used to measure how the structure of an RNA molecule changes as it emerges from an RNAP and has been applied to several riboswitches and non-coding RNAs^9,10,13,19–22^. While systematic cotranscriptional RNA structure probing experiments can detect structural rearrangements that occur as a nascent transcript grows longer, the ability of an intermediate structure to rearrange into another structure as transcription proceeds is not directly assessed by these methods because each reactivity profile is an end-point measurement. It is therefore possible that equilibration of a nascent transcript within a static TEC prior to chemical probing could cause the formation of a non-native structure that is not a true folding intermediate and which, in some cases, may not be able to rearrange into downstream native structures. This limitation of cotranscriptional RNA structure probing is partially addressed by the use of single-molecule force spectroscopy^12,18^ and single-molecule FRET^14–17^ to measure cotranscriptional RNA folding continuously with high temporal resolution. However, the ability to measure the cotranscriptional rearrangement of RNA structures by chemical probing would provide a means for assessing the validity of cotranscriptional RNA folding events that captures structural information for each nucleotide of the target RNA.

To facilitate the detection of cotranscriptional RNA folding events by chemical probing, we have developed linked-multipoint Transcription Elongation Complex RNA structure probing (TECprobe-LM). TECprobe-LM uses the SHAPE-MaP-based^23^ TECprobe platform^20^ that we developed previously to directly assess whether the reactivity profiles of two RNA folding intermediate populations are linked by a cotranscriptional folding event. In a TECprobe-LM experiment, aliquots of an *E. coli* RNAP *in vitro* transcription reaction are removed for chemical probing after RNAP has arrested at a photolabile NPOM-caged-dT^24^ stall site^25^ and after RNAP has arrested at a downstream biotin-streptavidin roadblock following removal of the NPOM-cage by irradiation with 365 nm UV light (Figure 1). In this way, the cotranscriptional conversion of one population of RNA structures into a second population of structures is directly measured by chemical probing. To validate this approach, we used TECprobe-LM to visualize the rearrangement of a non-native intermediate hairpin into the native *E. coli* signal recognition particle RNA, folding of the *C. beijerinckii pfl* ZTP riboswitch aptamer and expression platform, and folding of the *B. cereus crcB* fluoride riboswitch aptamer and expression platform. All cotranscriptional folding transitions were detected by TECprobe-LM, and the observed folding intermediates agreed with measurements made by systematic cotranscriptional RNA structure probing experiments in all but one case. The primary limitation of TECprobe-LM is that, like other cotranscriptional RNA structure probing methods, the representation of transcripts in a sequencing library can depend on the efficiency of an ssRNA ligation that is prone to sequence and structure biases. Nonetheless, it was possible to collect high-quality data for all samples described in this work, including one sample for which this ligation was particularly inefficient.

**Figure 1.**
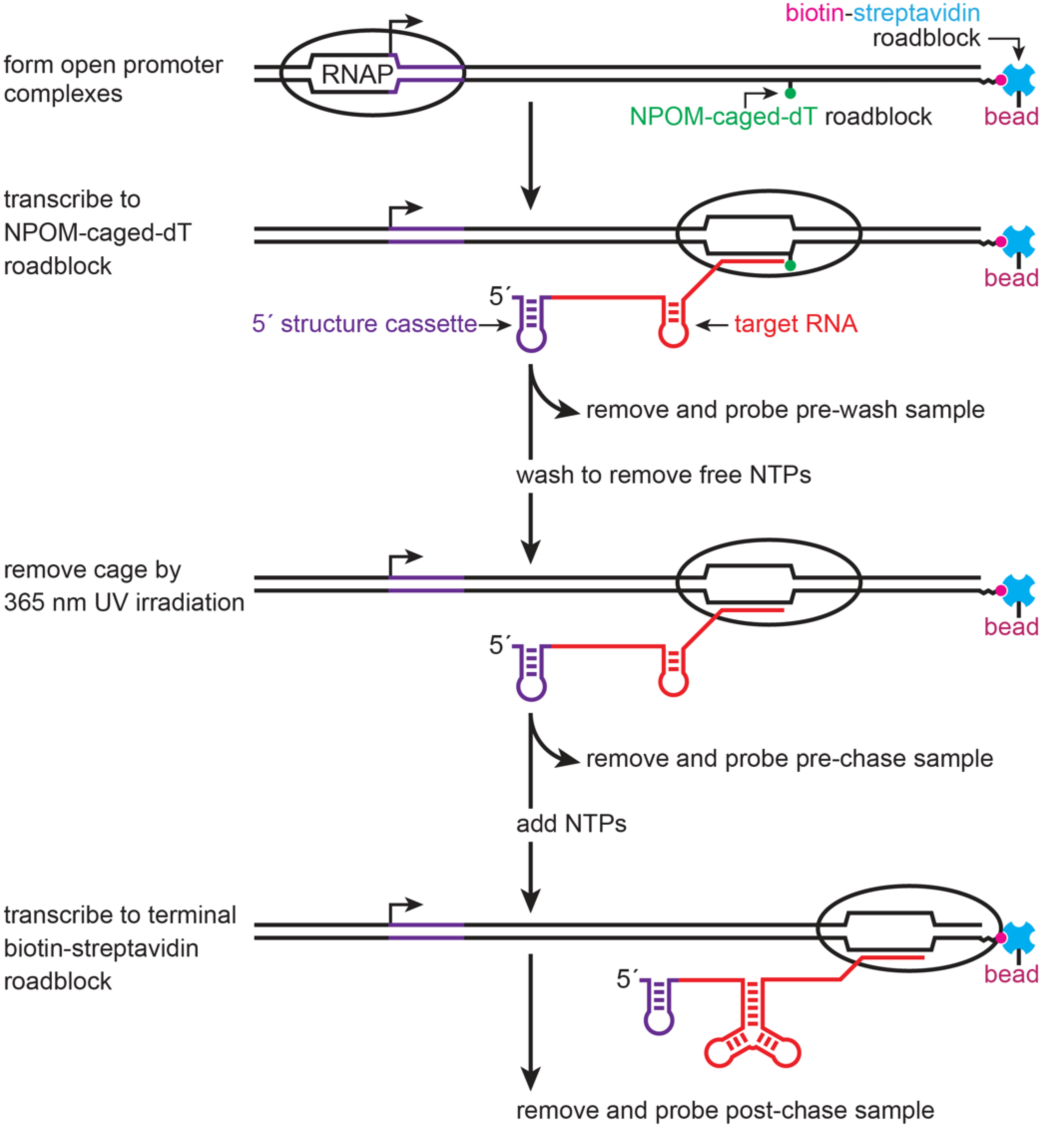
Overview of TECprobe-LM. Single-round transcription is initiated on template DNA that contains NPOM-caged-dT and terminal biotin-streptavidin roadblocks. In the initial phase of transcription, RNAP arrests at the NPOM-caged-dT roadblock. The pre-wash sample is removed from the reaction and chemically probed, and the remaining transcription reaction is washed to remove NTPs. The NPOM cage is removed by irradiation with 365 nm UV light and the pre-chase sample is removed from the reaction and chemically probed. Upon addition of NTPs, RNAP transcribes to the terminal biotin-streptavidin roadblock and the post-chase sample is removed and chemically probed. RNAP, RNA polymerase; SRP, signal recognition particle; SAv, streptavidin; BzCN, benzoyl cyanide.

Together, our findings establish TECprobe-LM as a strategy for measuring the cotranscriptional rearrangement of RNA structures using high-throughput RNA chemical probing.

## Results

### Overview of the linked multi-point cotranscriptional RNA structure probing strategy

Linked multi-point cotranscriptional RNA structure probing directly assesses whether one population of nascent RNA structures can cotranscriptionally rearrange into a second population of structures. This is accomplished by removing aliquots of an *E. coli* RNAP *in vitro* transcription reaction for chemical probing as RNAP is moved to specific locations within the DNA template (Figure 1). In the simplest implementation of TECprobe-LM described in this work, RNAP first transcribes to one nucleotide upstream of an NPOM-caged-dT modification in the template DNA strand, which was shown by Nadon et al. to function as a photoreversible transcription roadblock^25^. A ‘pre-wash’ aliquot of the transcription reaction is then removed for chemical probing and the arrested TECs are washed extensively to remove excess NTPs. The NPOM-photocage is then removed by exposure to 365 nm UV light and a ‘pre-chase’ aliquot of the transcription reaction is removed for chemical probing. NTPs are then added to the reaction so that RNAP transcribes to a terminal biotin-streptavidin roadblock and a ‘post-chase’ aliquot of the transcription reaction is removed for chemical probing. The pre-wash and pre-chase samples can be compared to ensure that the population of RNA structures that existed after RNAP first stalled at the NPOM-caged-dT site did not change when the arrested TECs were washed. The pre-chase and post-chase samples can be compared to assess how the population of RNA structures changed when RNAP transcribed downstream.

### Rearrangement of the 4.5S SRP RNA intermediate hairpin

Wong et al. previously showed that the *E. coli* signal recognition particle (SRP) RNA can form a non-native structure prior to folding of the native SRP RNA structure^8^. This model was later refined by Watters, Strobel et al. using Cotranscriptional SHAPE-Seq^9^. While these prior studies detected a non-native intermediate hairpin, they did not directly assess whether the intermediate hairpin could refold into the mature SRP RNA structure. Fukuda et al. later detected cotranscriptional refolding of the intermediate hairpin using single-molecule force spectroscopy^18^ and Yu et al. identified efficient mechanisms by which this structural transition could occur cotranscriptionally^19^. To visualize rearrangement of the SRP RNA intermediate hairpin by high-throughput RNA chemical probing, we performed a TECprobe-LM experiment in which RNAP was first positioned at +127, which precedes intermediate hairpin rearrangement, and then chased to +161 at which point the native SRP RNA structure is expected to have folded (Figure 2a).

**Figure 2.**
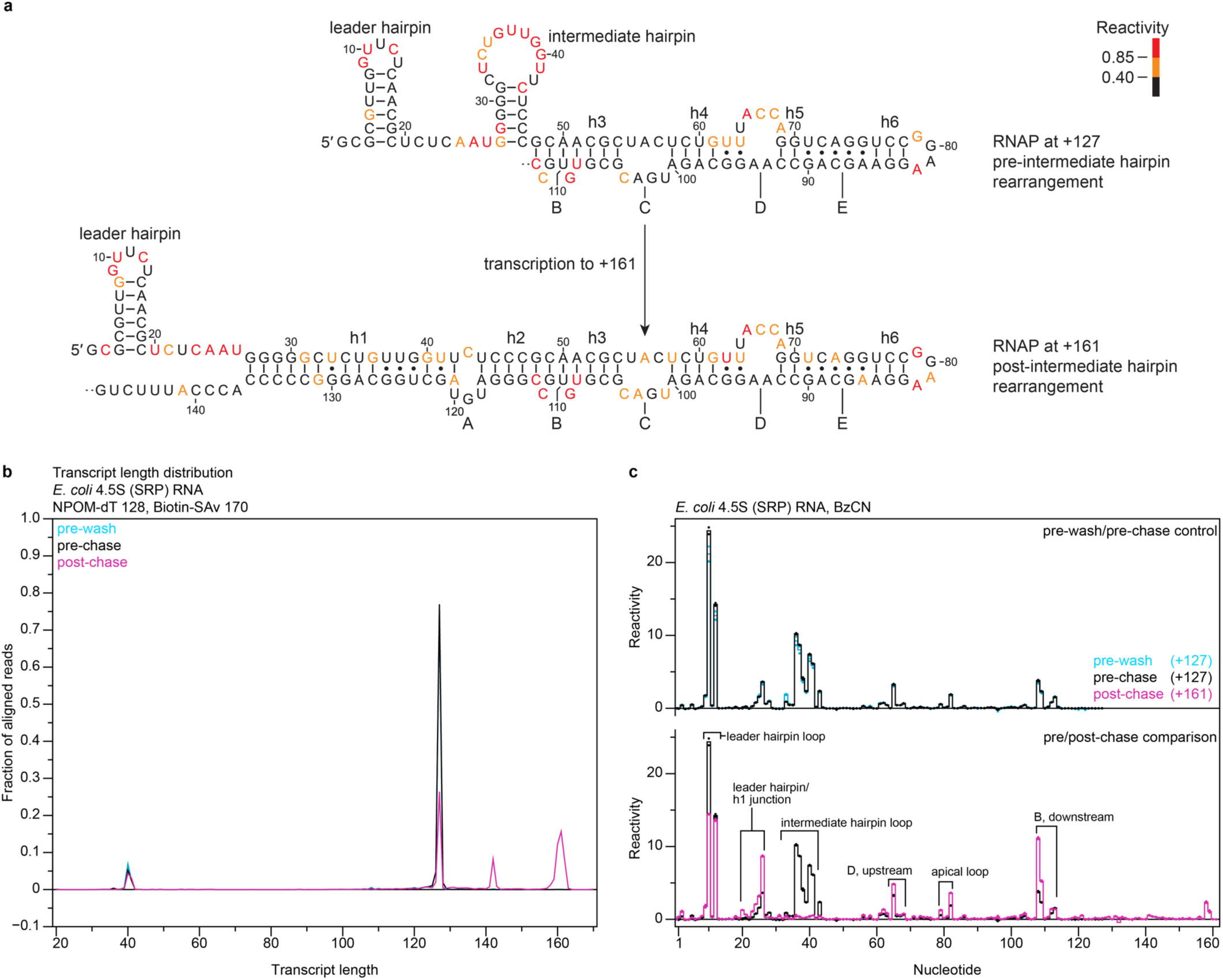
Cotranscriptional folding of the *E. coli* SRP RNA. (**a**) Secondary structures of the *E. coli* SRP RNA folding intermediates that were assessed by TECprobe-LM colored by reactivity. Sequence within the RNAP footprint is not shown. (**b**) Transcript length distribution for the pre-wash, pre-chase, and post-chase samples. Traces are the average of n=2 replicates. (**c**) Comparison of reactivity profiles for pre-wash and pre-chase samples (upper plot) and for pre-chase and post-chase samples (lower plot). Solid lines are the average of n=2 replicates and reactivity values for individual replicates are shown as points. RNAP, RNA polymerase; PK, pseudoknot; SAv, streptavidin; BzCN, benzoyl cyanide.

In the pre-wash and pre-chase samples, ∼98% of aligned reads mapped to transcripts upstream of the NPOM-caged-dT modification at +128, ∼75% mapped to +127, and ∼5% mapped to +126 (Figure 2b). In the post-chase sample, the fraction of aligned reads that mapped to transcripts beyond +127 increased from ∼2% to ∼57%, and ∼40% mapped to the biotin-streptavidin roadblock enrichment sites from +158 to +162 (Figure 2b). The presence of a roadblock-independent enrichment site at +142 suggests that RNAP is prone to arresting at this position (Figure 2b). Both the 142 nt transcript and biotin-streptavidin-enriched transcripts were observed by denaturing PAGE (Supplementary Figure 1a). However, in contrast to the read distribution observed by TECprobe-LM, the 142 nt transcript was more abundant than transcripts that were enriched by the biotin-streptavidin roadblock when assessed by gel electrophoresis. This difference is most likely caused by transcript-specific variation in the efficiency of 3’ adapter ligation skewing the representation of nascent transcripts in the sequencing library^13^.

Washing the roadblocked TECs to remove NTPs did not perturb RNA structure (Figure 2c, upper plot). Several known elements of the SRP RNA structure were observed when RNAP was positioned at +127: i) the 5’ leader hairpin and linker were detected as elevated reactivity at G9, U10, and C12 and at nts 24-26, respectively, ii) the intermediate hairpin loop was detected as elevated reactivity at nucleotides 33-41, and iii) the apical loop of the native SRP RNA structure, upstream segment of bulge D, and downstream segment of bulge B were detected as elevated reactivity at nucleotides within each of these regions of the SRP RNA hairpin (Figure 2a, c, lower plot). Upon transcription from +127 to +161, nucleotides within the intermediate hairpin loop become non-reactive while the reactivity of flexible nucleotides in the native SRP RNA structure and the 5’ leader persists, indicating that the non-native intermediate hairpin has refolded (Figure 2a, c, lower plot). The reactivity profiles obtained using TECprobe-LM agreed with end-point profiles collected using variable length TECprobe (TECprobe-VL) except that the leader hairpin and intermediate hairpin loops were more reactive in TECprobe-LM experiments (Supplementary Figure 2).

### *C. beijerinckii pfl* ZTP riboswitch aptamer folding

The ZTP aptamer comprises two sub-domains that form a long-range pseudoknot^26^ (Figure 3a, 120 nt transcript). ZMP binding establishes a contiguous helical stack between P3 and the pseudoknot (PK) which, in the *C*. *beijerinckii pfl* ZTP aptamer, blocks nucleation of the terminator hairpin^11,27–30^. To visualize *pfl* aptamer folding, we initially performed a TECprobe-LM experiment in which RNAP was first positioned at +102, which precedes pseudoknot and P3 subdomain folding, and then chased to +120 at which point the pseudoknot and P3 are folded and ZMP can bind.

**Figure 3.**
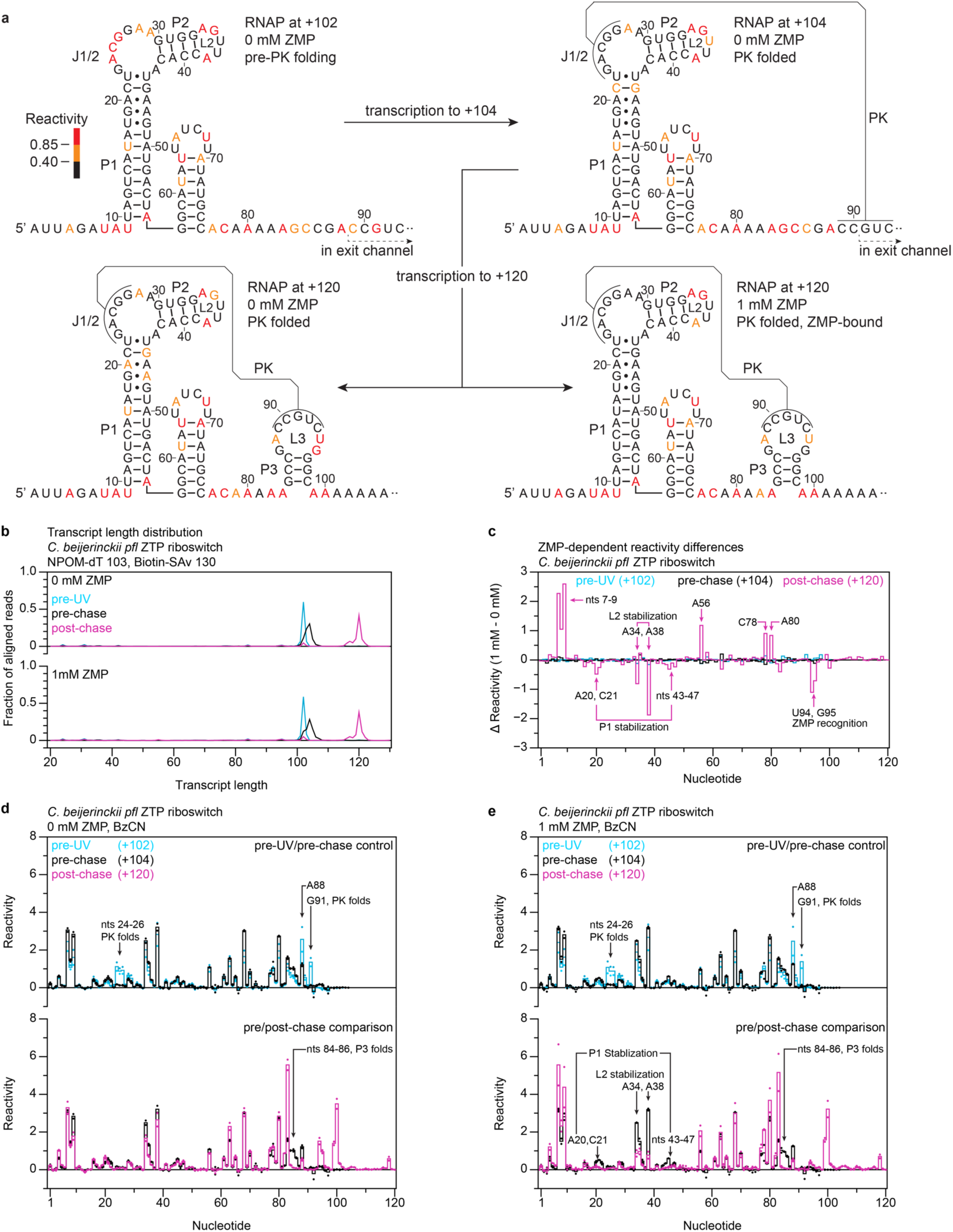
Cotranscriptional folding of the *C*. *beijerinckii pfl* ZTP riboswitch aptamer. (**a**) Secondary structures of the *C. beijerinckii pfl* ZTP riboswitch aptamer folding intermediates that were assessed by TECprobe-LM colored by reactivity. (**b**) Transcript length distribution for the pre-UV, pre-chase, and post-chase samples. Traces are the average of n=2 replicates. (**c**) Difference in reactivity (1 mM ZMP - 0 mM ZMP) observed for the pre-UV, pre-chase, and post-chase samples. Differences were calculated from the average reactivity values shown in (d) and (e). (**d**, **e**) Comparison of reactivity profiles for pre-UV and pre-chase samples (upper plot) and for pre-chase and post-chase samples (lower plot) in the absence (d) and presence (e) of 1 mM ZMP. Solid lines are the average of n=2 replicates and reactivity values for individual replicates are shown as points. RNAP, RNA polymerase; PK, pseudoknot; SAv, streptavidin; BzCN, benzoyl cyanide.

In the pre-wash sample, ∼91% of aligned reads mapped to transcripts upstream of the NPOM-caged-dT modification at +103, ∼60% mapped to +102, and ∼4% mapped to +101 (Supplementary Figure 3a). ∼6% of aligned reads mapped to +103, which indicates that RNAP can insert a nucleotide opposite of the NPOM-caged-dT modification in some sequence contexts including the poly-dT tract of the current construct. In the pre-chase sample, RNAP was able to incorporate 1-2 nucleotides following UV irradiation despite extensive washing to remove excess NTPs (Supplementary Figure 3a). Comparison of reactivity profiles for these transcripts revealed that translocation of RNAP from +102 to +103 permits the ZTP aptamer pseudoknot to fold (Supplementary Figure 3b). Consequently, TECs in the pre-chase sample comprise a mixed population in which ≥103 nt RNAs have formed the pseudoknot and 102 nt RNAs have not. In the post-chase sample, the fraction of aligned reads that mapped to transcripts beyond +103 increased from ∼2% to >80-90%, and ∼80- 84% mapped to the biotin-streptavidin roadblock enrichment sites from +116 to +122 (Supplementary Figure 3a). The transcript distribution observed by sequencing agreed with the distribution observed by denaturing PAGE (Supplementary Figure 1b).

We reasoned that one way to resolve the mixed population that was observed in the pre-chase sample is to favor translocation to ≥+103 by increasing the concentration of NTPs used during the initial phase of transcription and washing the roadblocked TECs less extensively. This would enable a three-point TECprobe- LM experiment in which two sequential cotranscriptional RNA folding transitions are observed (Figure 3a). In this format, the first sample that is collected is referred to as the ‘pre-UV’ sample, and the pre-wash control is omitted for simplicity because washing TECs to remove NTPs did not perturb RNA structure when the experiment was performed using the standard TECprobe-LM format (Supplementary Figure 3c). Increasing NTP concentration and reducing the wash volume caused ∼86% of TECs that were arrested at the NPOM-caged-dT stall site to transcribe to positions +103 to +106 upon release of the NPOM cage (Figure 3b). The reactivity profiles of the resulting 103 to 106 nt transcripts are virtually identical (Supplementary Figure 3d). As observed in the initial experiment described above, ∼80% of aligned reads mapped to the biotin-streptavidin enrichment sites from +116 to +122 in the post-chase sample (Figure 3b).

As expected, ZMP binding was not detected when RNAP was positioned at +102 because the aptamer has not yet folded (Figure 3c). In both the absence and presence of ZMP, transcription to +104 caused the reactivity of nucleotides 24-26, A88, and G91 to decrease upon pseudoknot folding (Figure 3a, d, e, upper plots). ZMP binding was still not detected because the P3 hairpin has not yet folded (Figure 3c). Chasing RNAP to +120 in the absence of ZMP caused the reactivity of nucleotides 84-86 to decrease as P3 folds (Figure 3a, d, lower plot). When RNAP was chased to +120 in the presence of ZMP, P3 folding was observed in coordination with expected ZMP-dependent reactivity changes including: i) decreased reactivity in P1 due to stabilization of non-canonical base pairs, ii) decreased reactivity of A34 and A38 in L2 which corresponds to formation of a conserved A-minor motif, and iii) decreased reactivity at U94 and G95 in L3 which hydrogen bond and stack with the Z nucleobase, respectively^11,20^ (Figure 3a, c, e, lower plot). The reactivity profiles obtained using TECprobe-LM agree with profiles collected using TECprobe-VL with two exceptions: First, the reactivity of nucleotides immediately upstream of the RNAP footprint in the 102 nt transcript is higher in the TECprobe-LM profile (Supplementary Figure 4a, b). Second, the TECprobe-LM profiles of the 104 nt transcript more closely match the TECprobe-VL profiles for the 110 nt transcript, in which the pseudoknot has fully folded, than the TECprobe-VL profiles for the 104 nt transcript (Supplementary Figure 4). Both exceptions are most likely caused by backtracking that can occur when RNAP collides with a biotin-streptavidin roadblock during the TECprobe-VL procedure^13^, which shifts the RNAP footprint upstream and prevents the formation of RNA structures that would otherwise fold if the nascent transcript 3’ end were positioned at the RNAP active center.

### *C. beijerinckii pfl* ZTP riboswitch terminator folding

To visualize *pfl* ZTP riboswitch terminator hairpin folding, we performed a TECprobe-LM experiment in which RNAP was first positioned at +111, at which point P3 can fold while partially in the RNA exit channel of RNAP, and then chased downstream of the termination site to +143 (Figure 4a, b). In this experimental configuration, ZMP binding occurs cotranscriptionally as the ZTP aptamer emerges from RNAP until ∼+123 when the terminator hairpin can nucleate.

**Figure 4.**
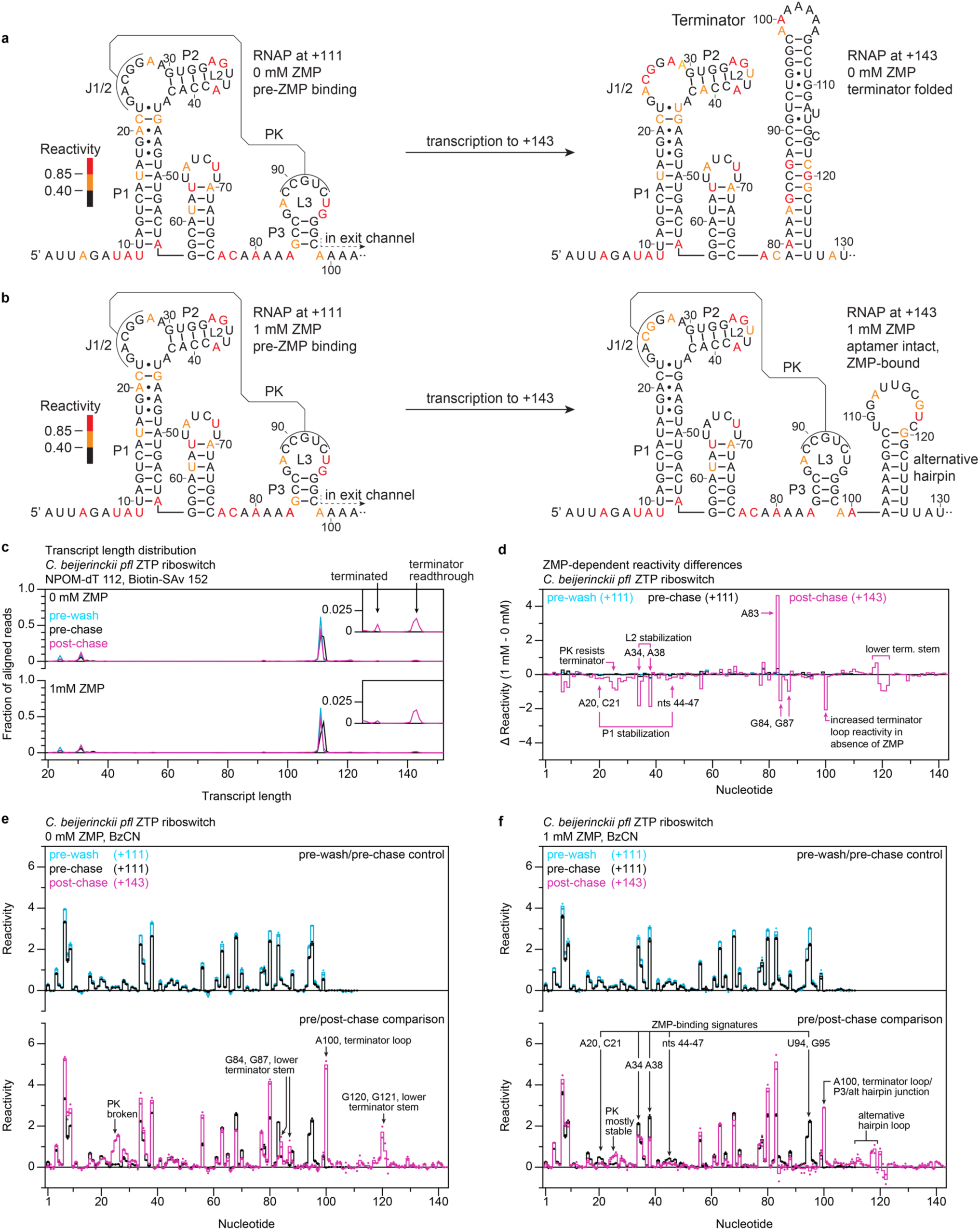
Cotranscriptional folding of the *C. beijerinckii pfl* ZTP riboswitch expression platform. (**a**, **b**) Secondary structures of the *C. beijerinckii pfl* riboswitch expression platform folding intermediates that were assessed by TECprobe-LM in the absence (a) and presence (b) of 1 mM ZMP colored by reactivity. (**c**) Transcript length distribution for the pre-wash, pre-chase, and post-chase samples. Traces are the average of n=2 replicates. (**d**) Difference in reactivity (1 mM ZMP - 0 mM ZMP) observed for the pre-wash, pre-chase, and post-chase samples. Differences were calculated from the average reactivity values shown in (e) and (f). (**e**, **f**) Comparison of reactivity profiles for pre-wash and pre-chase samples (upper plot) and for pre-chase and post-chase samples (lower plot) in the absence (e) and presence (f) of 1 mM ZMP. Solid lines are the average of n=2 replicates and reactivity values for individual replicates are shown as points. RNAP, RNA polymerase; PK, pseudoknot; SAv, streptavidin; BzCN, benzoyl cyanide.

In the pre-wash sample, ∼97% of aligned reads mapped to transcripts upstream of the NPOM-caged-dT modification at +112, ∼62% mapped to +111, and ∼4% mapped to +110 (Figure 4c). In the pre-chase sample, NPOM-caged-dT-enriched transcripts were observed at +111 and +112, which indicates that RNAP was able to incorporate one additional nucleotide upon release of the NPOM cage in some cases despite extensive washing (Figure 4c). The reactivity profiles of the 111 nt and 112 nt transcripts were indistinguishable in the absence of ZMP (Supplementary Figure 5a, upper plot). In the presence of ZMP, the reactivity of ZMP-responsive nucleotides in the 112 nt transcript decreased to a value between that of the apo aptamer and the ZMP-bound aptamer (Supplementary Figure 5a, lower plot, 5b). This indicates that 112 nt transcripts can bind ZMP to some extent, although it is not clear whether the intermediate reactivity values are caused by binding of ZMP to a subset of aptamers or by interconversion between apo and ZMP-bound states. Denaturing PAGE analysis of post-chase transcripts shows that most TECs that are positioned at +111/+112 resume transcription and yield terminated and full-length RNA products (Supplementary Figure 1c). In contrast with this observation, <5% of reads mapped to transcripts at the termination site or biotin-streptavidin roadblock in the post-chase sample (Figure 4c). The depletion of terminated and full-length transcripts in TECprobe-LM libraries is likely caused by inefficient 3’ adapter ligation and/or reverse transcription and was observed previously in cotranscriptional SHAPE-Seq and TECprobe-VL libraries^13,20^. Despite this issue, the sequencing depth of full-length transcripts was sufficient to generate high-quality reactivity profiles and it was possible to assess expression platform folding by comparing the reactivity profiles of transcripts within TECs that were arrested at +111/+112 to transcripts within TECs that were arrested at +143.

Washing the roadblocked TECs to remove NTPs did not perturb RNA structure (Figure 4e, f, upper plots). As expected, ZMP binding was not detected when RNAP was positioned at +111 because the aptamer has not yet fully emerged from RNAP (Figure 4d). As noted above, the pre-chase sample also contained TECs at +112 for which partial ZMP binding was observed (Supplementary Figure 5). Terminator folding was observed when RNAP was chased to +143 in the absence of ZMP as: i) increased reactivity at nucleotides 24-26 indicating that the pseudoknot was broken, ii) increased reactivity at A100 within the terminator loop, which reflects the reactivity of the terminator loop in general due to the ambiguity of mapping mutations within a homopolymer, and iii) decreased reactivity in the terminator stem except for nucleotides within the lower segment of the stem, all of which are consistent with previous TECprobe-VL measurements^20^ (Figure 4a, e, lower plot). When RNAP was chased to +143 in the presence of 1 mM ZMP, the pseudoknot remained mostly intact, all signatures of ZMP binding described above for the ZTP aptamer folding transition were detected, and reactivity signatures consistent with a proposed alternative structure that forms upon transcription antitermination were detected (Figure 4b, d, f, lower plot). This indicates that ZMP binding caused a detectable fraction of ZTP riboswitches to antiterminate transcription. The reactivity profiles measured by TECprobe-LM agreed with profiles measured by TECprobe-VL except that the reactivity of nucleotides immediately upstream of the RNAP footprint in the 111 nt transcript was higher in the TECprobe-LM data (Supplementary Figure 6). As above, this is most likely explained by biotin-streptavidin roadblock-induced backtracking^13^ that does not occur when RNAP is arrested at an NPOM-caged-dT modification.

### *B. cereus crcB* fluoride riboswitch aptamer folding

The fluoride aptamer comprises an H-type pseudoknot and several long-range base pairs that facilitate the coordination of fluoride by three Mg^2+^ ions^31,32^ (Figure 5a, b, 71 nt transcript). Fluoride binding locks the aptamer into a conformation in which the pseudoknot and the A40:U48 linchpin base pair are stably formed that, in the case of the *B. cereus crcB* fluoride riboswitch, causes transcription antitermination^9,33,34^. To visualize *crcB* fluoride aptamer folding, we performed a TECprobe-LM experiment in which RNAP was first positioned at +54, which precedes aptamer folding, and then chased to +71 at which point the pseudoknot is stably folded and the aptamer is competent to bind fluoride (Figure 5a, b).

**Figure 5.**
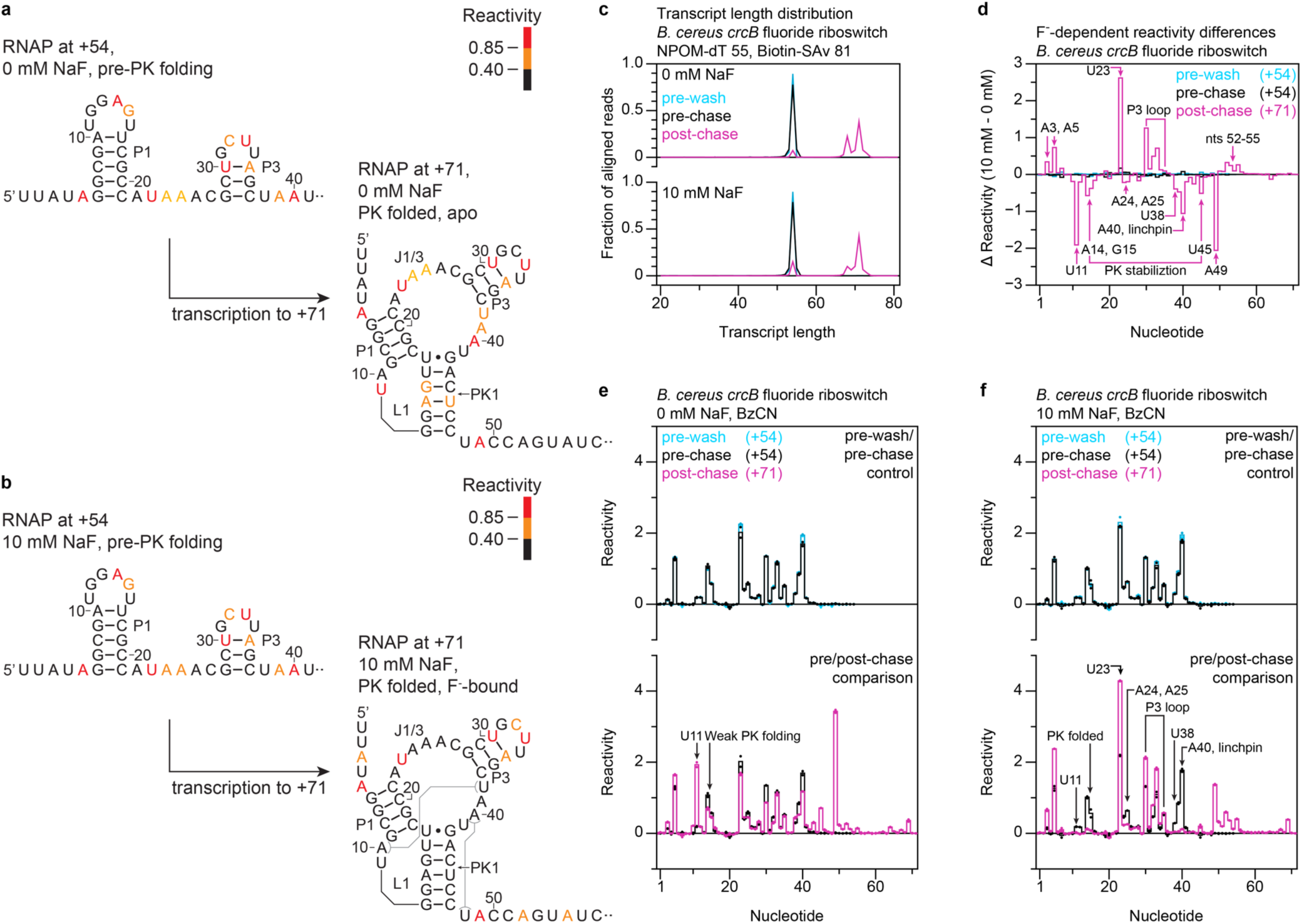
Cotranscriptional folding of the *B. cereus crcB* fluoride riboswitch aptamer. (**a**, **b**) Secondary structures of the *B. cereus crcB* fluoride riboswitch aptamer folding intermediates that were assessed by TECprobe-LM in the absence (a) and presence (b) of 10 mM NaF colored by reactivity. Sequence within the RNAP footprint is not shown. (**c**) Transcript length distribution for the pre-wash, pre-chase, and post-chase samples. Traces are the average of n=2 replicates. (**d**) Difference in reactivity (10 mM NaF - 0 mM NaF) observed for the pre-wash, pre-chase, and post-chase samples. Differences were calculated from the average reactivity values shown in (e) and (f). (**e**, **f**) Comparison of reactivity profiles for pre-wash and pre-chase samples (upper plot) and for pre-chase and post-chase samples (lower plot) in the absence (e) and presence (f) of 10 mM NaF. Solid lines are the average of n=2 replicates and reactivity values for individual replicates are shown as points. RNAP, RNA polymerase; PK, pseudoknot; SAv, streptavidin; BzCN, benzoyl cyanide.

In the pre-wash samples, ∼99% of aligned reads mapped to transcripts upstream of the NPOM-caged-dT modification at +55, ∼89% mapped to +54, and ∼5% mapped to +53 (Figure 5c). In the pre-chase samples, ∼78% of aligned reads mapped to +54, ∼8% mapped to +53, and ∼8% mapped to +55 (Figure 5c). In the post-chase samples, >80% of aligned reads mapped to the biotin-streptavidin roadblock enrichment sites from +67 to +73 (Figure 5c). The transcript distribution observed by sequencing agreed with the distribution observed by denaturing PAGE (Supplementary Figure 1d).

Washing the roadblocked TECs to remove NTPs did not perturb RNA structure (Figure 5e, f, upper plots). As expected, fluoride had no effect on RNA structure when RNAP was positioned at +54 because the aptamer has not yet folded (Figure 5d). Chasing RNAP from +54 to +71 in the absence of fluoride caused an increase in the reactivity of U11, which caps an extended P3 stack that is stabilized by fluoride binding^32^ (Figure 5a, e, lower plot). In previous TECprobe-VL experiments that were performed without fluoride, increased U11 reactivity was observed in coordination with decreased reactivity at A14 and G15 due to pseudoknot folding^20^ (Supplementary Figure 7a, b, upper plots). This implies that U11 becomes reactive as other nucleotides within L1 form pseudoknot base pairs in the absence of fluoride. However, in the current TECprobe-LM data, the decrease in reactivity at A14 and G15 when RNAP transcribes from +54 to +71 is negligible (Figure 5e, lower plot). It is unlikely that the pseudoknot can fold when RNAP is positioned at +54 because two of six downstream pseudoknot nucleotides are expected to be paired within the RNA-DNA hybrid. Furthermore, the reactivity profiles of the 53 and 52 nt transcripts, in which additional downstream pseudoknot nucleotides are paired within the RNA-DNA hybrid, are identical to that of the 54 nt transcript except that A40 is protected from benzoyl cyanide modification when RNAP is at +52 (Supplementary Figure 8). This indicates that the approximately constant reactivity observed at A14 and G15 is not due to the pseudoknot folding at +54 and that fluoride-independent pseudoknot folding is not detected in the TECprobe-LM experiment. Comparison of TECprobe-LM and TECprobe-VL reactivity profiles for the 54 nt transcript indicates that the nascent RNA adopts a distinct conformation in each experiment (Supplementary Figure 7a, c, upper plots). Most notably, nucleotides A14 and G15 within L1 and G28, U30 and A35 in P3 are more reactive in the TECprobe-VL profile than in the TECprobe-LM profile. The latter observation suggests that P3 has stably folded when RNAP is positioned at +54 in the TECprobe-LM experiment but not in the TECprobe-VL experiment. In further support of this interpretation, the TECprobe-LM profile for the 54 nt transcript is similar to the TECprobe-VL profile for the 68 nt transcript, except that U11 is reactive at +68 but not at +54 (Supplementary Figure 7b, upper plot).

This indicates that during the TECprobe-LM experiment, P3 folds earlier than was observed by TECprobe-VL. While the cause of this difference is unclear, the agreement of the TECprobe-LM and TECprobe-VL reactivity profiles for the 71 nt transcript in both ligand conditions indicates that the reactivity differences observed at +54 do not interfere with aptamer folding or fluoride binding (Supplementary Figure 7a, c, lower plots). The observation that transcription from +54 to +71 causes U11 to become reactive in the absence of fluoride and causes fluoride-dependent stabilization of the pseudoknot in the presence of fluoride indicates that the pseudoknot likely folds transiently until fluoride binds. This is consistent with observations made by single-molecule FRET, in which the *crcB* fluoride aptamer exists in a dynamic docked state until fluoride binding stabilizes the pseudoknot^34^.

Established fluoride-dependent reactivity changes were observed when RNAP was chased from +54 to +71 in the presence of fluoride, including: i) decreased reactivity at A14, G15, and U45 due to pseudoknot stabilization, ii) decreased reactivity at U38, which forms a reversed Watson-Crick pair with A10 that extends the P3 stack, and at U11, which caps the extended P3 stack, iii) decreased reactivity at A40, which forms the linchpin base pair with U48, iv) decreased reactivity at A49, which stacks with the linchpin base pair, v) increased reactivity at U23 and decreased reactivity at A24 and A25 within J1/3, and vi) increased reactivity at U30, C32, and U33 within the P3 stem and loop^20^ (Figure 5b, d, f, lower plot). The reactivity profiles for the 71 nt transcript obtained using TECprobe-LM agreed with the TECprobe-VL profiles except that the reactivity of nucleotides immediately upstream of the RNAP footprint was lower in the TECprobe-LM data (Supplementary Figure 7a, c, lower plots). This difference is most likely caused by the terminal biotin-streptavidin roadblock inducing more extensive backtracking in the TECprobe-LM experiment than the internal biotin-streptavidin roadblocks in the TECprobe-VL experiment.

### *B. cereus crcB* fluoride riboswitch terminator folding

To visualize *crcB* fluoride riboswitch terminator hairpin folding, we performed a TECprobe-LM experiment in which RNAP was first positioned at +74, after the fluoride aptamer has folded and fluoride can bind, and then chased downstream of the termination site to +95 (Figure 6a, b). In the pre-wash sample, 99% of aligned reads mapped to transcripts upstream of the NPOM-caged-dT modification at +75, ∼88% mapped to +74, and ∼5% mapped to +73 (Figure 6c). In the pre-chase sample, ∼70% of aligned reads mapped to +74 and ∼16% mapped to +73 (Figure 6c). In the post-chase sample, ∼71-76% of aligned reads mapped to the biotin-streptavidin roadblock enrichment sites from +93 to +96 (Figure 6c). Although both terminated and full-length transcripts were detected by denaturing PAGE, terminated transcripts were depleted in the TECprobe-LM sequencing library (Figure 6c, Supplementary Figure 1e). This is most likely caused by inefficient ligation of the 3’ adapter since full-length transcripts face the same reverse transcription barriers as terminated transcripts. Nonetheless, it was possible to assess expression platform folding by comparing the reactivity profiles of transcripts within TECs that were arrested at +74 to transcripts within TECs that were arrested at +95.

**Figure 6.**
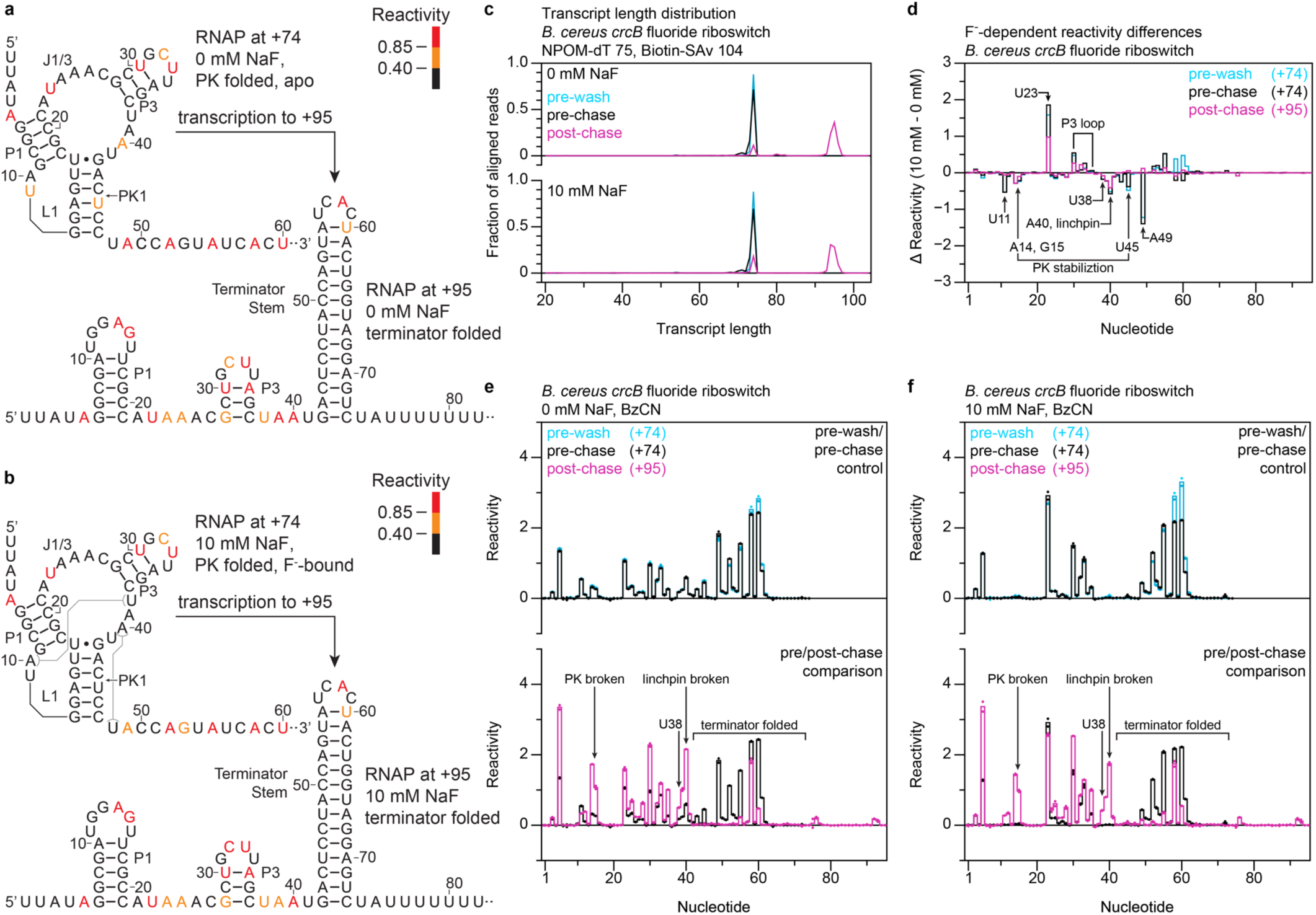
Cotranscriptional folding of the *B. cereus crcB* fluoride riboswitch expression platform. (**a**, **b**) Secondary structures of the *B. cereus crcB* fluoride riboswitch expression platform folding intermediates that were assessed by TECprobe-LM in the absence (a) and presence (b) of 10 mM NaF colored by reactivity. Sequence within the RNAP footprint is not shown. (**c**) Transcript length distribution for the pre-wash, pre-chase, and post-chase samples. Traces are the average of n=2 replicates. (**d**) Difference in reactivity (10 mM NaF - 0 mM NaF) observed for the pre-wash, pre-chase, and post-chase samples. Differences were calculated from the average reactivity values shown in (e) and (f). (**e**, **f**) Comparison of reactivity profiles for pre-wash and pre-chase samples (upper plot) and for pre-chase and post-chase samples (lower plot) in the absence (e) and presence (f) of 10 mM NaF. Solid lines are the average of n=2 replicates and reactivity values for individual replicates are shown as points. RNAP, RNA polymerase; PK, pseudoknot; SAv, streptavidin; BzCN, benzoyl cyanide.

Washing the roadblocked TECs to remove NTPs did not perturb the structure of the fluoride aptamer but caused a small decrease in the reactivity of nucleotides A58, U60, and A61 that was larger in the presence of fluoride (Figure 6e, f, upper plots). As expected, fluoride binding was detected when RNAP was positioned at +74 because the fluoride aptamer had folded (Figure 6d). In both the absence and presence of 10 mM fluoride, chasing RNAP to +95 caused known signatures of terminator hairpin folding^20^ including: i) increased reactivity at A14 and G15 due to pseudoknot disruption, ii) increased reactivity at nucleotides 38-40 due to disruption of the A10:U38 and A40:U48 pairs, and iii) decreased reactivity in terminator stem nucleotides (Figure 6a, b, e, f, lower plots). The partial persistence of fluoride-dependent differences between the 0 mM and 10 mM NaF post-chase samples indicates that some fluoride aptamers did remain intact in the presence of the terminator hairpin (Figure 6d).

The TECprobe-LM reactivity profiles for the 74 nt transcript agreed with the TECprobe-VL profiles except that the reactivity of nucleotides immediately upstream of the RNAP footprint was higher in the TECprobe-LM data (Supplementary Figure 9a, c, upper plots). As described above, this is most likely caused by biotin-streptavidin roadblock-induced backtracking shifting the RNAP footprint upstream in the TECprobe-VL experiment^13^. The TECprobe-LM and TECprobe-VL profiles for transcript 95 agreed except that terminator hairpin stem reactivity was lower in the 0 mM NaF TECprobe-LM data, which may be due to ∼82-fold higher sequencing depth yielding a higher quality reactivity profile (Supplementary Figure 9a, c, lower plots). In support of this interpretation, terminator stem nucleotides are weakly reactive in the TECprobe-VL profile of the 80 nt terminated transcript, which was sequenced ∼42.5-times deeper than the 95 nt transcript (Supplementary Figure 9b).

## Discussion

TECprobe-LM uses RNA chemical probing to directly assess whether one RNA folding intermediate can cotranscriptionally rearrange into another folding intermediate. The primary advance of the linked-multipoint cotranscriptional RNA structure probing strategy relative to previous approaches is that it assesses whether two RNA folding intermediates are linked by a cotranscriptional RNA folding event, which was not previously possible to measure by chemical probing. However, like other cotranscriptional RNA structure probing methods, there is a possibility that nascent RNA structures will equilibrate after transcription has been arrested. Because of this, the linked-multipoint strategy for cotranscriptional RNA chemical probing does not measure true cotranscriptional RNA folding like single-molecule force spectroscopy^12,18^ and single-molecule FRET experiments^14–17^. Nonetheless, the ability to assess whether one population of nascent RNA structures can rearrange into another population of structures using high-throughput RNA structure probing is complementary to these biophysical approaches because it yields structural information for each nucleotide of the transcript.

There are several technical limitations that must be considered when designing linked-multipoint cotranscriptional RNA structure probing experiments. First, although many photocaged nucleotides have been incorporated into DNA^35,36^, only NPOM-caged-dT is currently commercially available. For most users, this limits the potential sites that can be used to reversibly halt RNAP. Second, like other cotranscriptional RNA structure probing methods, TECprobe-LM requires the ligation of a sequencing adapter to the nascent RNA 3’ end. As described previously, the efficiency of this ligation can vary depending on RNA sequence and structure^13,37,38^. Consequently, the distribution of transcript lengths that is generated by the transcription reaction may not be accurately captured within a sequencing library. The effects of these biases were observed in several ways in the systems assessed here. For example, terminated fluoride riboswitch transcripts were nearly undetectable in TECprobe-LM data but were clearly observed by gel electrophoresis. Similarly, terminated and full-length ZTP riboswitch transcripts were severely underrepresented in TECprobe-LM data but were clearly observed by gel electrophoresis. This limitation cannot currently be circumvented but also did not prevent high-quality chemical probing data from being obtained in the systems that were assessed in this work. Third, the sequence context of the NPOM-caged-dT modification can affect experiment outcomes. For example, TECs that were arrested at the NPOM-caged-dT modification in the SRP RNA DNA template resumed transcription less efficiently than the TECs that were arrested at other NPOM-caged-dT sites. For these reasons, the success of a TECprobe-LM experiment is dependent on the target RNA sequence.

In general, the reactivity profiles obtained using TECprobe-LM agreed with end-point cotranscriptional RNA structure probing measurements made by TECprobe-VL. However, in several samples the reactivity of nucleotides that would be immediately outside of the RNA exit channel if the RNA 3’ end were positioned at the RNAP active center was higher when RNAP was arrested at an NPOM-caged-dT roadblock. This observation is consistent with the prior observation that collision of an *E. coli* TEC with a biotin-streptavidin complex can cause RNAP to backtrack, which would shift the RNAP footprint upstream^13^. For this reason, it is critical to consider the uncertainty in when a folding transition occurs when using TECprobe-VL data to select sites at which RNAP will be arrested by an NPOM-caged-dT modification or any other chemically encoded transcription roadblock. More notably, in the TECprobe-LM reactivity profile of the 54 nt fluoride riboswitch transcript, P3 folded earlier than in the TECprobe-VL experiment and the reactivity of L1 did not change upon pseudoknot folding. While the cause of this difference is not clear, notable differences in the procedures include the use of solution (VL) vs solid-phase (LM) transcription, and the inclusion of free streptavidin in the transcription reaction (VL). Nonetheless, the TECprobe-LM and TECprobe-VL datasets converged upon the same structure once RNAP transcribed downstream to +71 despite having different starting structures. We anticipate that TECprobe-VL data will typically be used to inform the design of TECprobe-LM experiments, which will facilitate comparisons between measurements made by the two methods as we have presented here.

The basic TECprobe-LM procedure is designed to assess one cotranscriptional RNA folding event at a time. Nonetheless, it was possible to assess two sequential *pfl* ZTP aptamer folding events by adjusting NTP concentration and depletion conditions so that RNAP transcribed forward several nucleotides upon release of the NPOM-cage. This approach took advantage of the observation that RNAP was prone to transcribing 1-2 nucleotides forward in this sequence context even after extensive washing to deplete NTPs and may not be easily generalizable. The simplest way to implement a TECprobe-LM experiment for >2 folding intermediates is to position RNAP at an initial NPOM-caged-dT modification and subsequently walk RNAP to specific downstream sites by resuming transcription in the absence of one or more NTPs. However, the complexity of a TECprobe-LM experiment increases substantially as more folding intermediates are assessed both because the efficiency of walking RNAP is sequence-dependent and because TECs that do not resume transcription after an initial walk may resume transcription during a subsequent walk. Furthermore, every additional intermediate that is assessed in a TECprobe-LM experiment requires two or four additional samples depending on whether a pre-wash control sample is collected. For these reasons, we anticipate that the most straightforward way to assess the validity of a sequence of RNA folding transitions will be to perform several independent TECprobe-LM experiments that assess overlapping folding transitions as we have done in the ZTP and fluoride riboswitch systems.

## Online Methods

### Oligonucleotides

All oligonucleotides were purchased from Integrated DNA Technologies. A detailed description of all oligonucleotides including sequence, modifications, and purifications is presented in Supplementary Table 1.

### Proteins

Q5 High-Fidelity DNA Polymerase, Vent (exo-) DNA polymerase, *Sulfolobus* DNA Polymerase IV, Lambda Exonuclease, *E*. *coli* RNA Polymerase holoenzyme, Mth RNA Ligase (as part of the 5’ DNA Adenylation kit), T4 RNA Ligase 2 truncated KQ, ET SSB, RNase H, and RNase I_f_ were purchased from New England Biolabs. TURBO DNase, SuperaseIN, SuperScript II, and BSA were purchased from ThermoFisher. Streptavidin was purchased from Promega.

### DNA template purification

DNA templates that contained an internal NPOM-caged-dT and 5’ biotin modification were prepared under 592 nm amber light by one of two strategies, which are described in detail below. In the first strategy, PCR amplification was performed using a primer that contained the NPOM-caged-dT and 5’ biotin modifications.

Translesion DNA synthesis was then performed to fill in the 5’ overhang that results from the NPOM-caged-dT modification blocking complete synthesis of the non-transcribed DNA strand. We previously used this strategy to synthesize DNA templates that contain internal biotin-TEG^39^, desthiobiotin-TEG, etheno-dA, and amino linker modifications^40^, however the success of translesion synthesis past the NPOM-caged-dT modification was variable depending on DNA template sequence. In the second strategy, we circumvented this issue by performing the initial PCR amplification using a 5’-phosphorylated reverse primer so that the transcribed DNA strand could be selectively degraded by lambda exonuclease and resynthesized using a primer that contained the NPOM-caged-dT and 5’ biotin modifications.

In all DNA template preparations, an unmodified linear dsDNA template was first amplified from plasmid DNA. PCR was performed as three 100 μl reactions containing 1X Q5 Reaction Buffer, 200 μM dNTPs, 250 nM PRA1_NoMod.F (Supplementary Table 1), 250 nM of a reverse primer that varied depending on the target sequence and the DNA template preparation strategy (Supplementary Tables 1 and 2), 0.2 ng/μl plasmid DNA, and 0.02 U/μl Q5 DNA polymerase using the following thermal cycler protocol with a heated lid set to 105 °C: 98 °C for 30 s, [98 °C for 10 s, 65 °C for 30 s, 72 °C for 20 s] x 30 cycles, 72 °C for 2 min, hold at 12 °C. For DNA template preparations in which the NPOM-caged-dT was added by PCR and translesion DNA synthesis, the 5’ end of the reverse primer was either positioned 1 nt before the position at which the NPOM-caged-dT modification would be located in the final dsDNA template or the primer overlapped this position. For DNA template preparations in which the NPOM-caged-dT modification was added by lambda exonuclease treatment and primer extension, the reverse primer matched the sequence of the NPOM-caged-dT-modified reverse primer that would be used for primer extension, but did not contain the NPOM-caged-dT modification and was 5’-phosphorylated. Linear dsDNA was then ethanol precipitated, purified by UV-free agarose gel extraction using a QIAquick gel extraction kit, and quantified using the Qubit dsDNA Broad Range Assay kit with a Qubit 4 Fluorometer exactly as described previously^39^.

When preparing DNA templates by PCR and translesion DNA synthesis, the NPOM-caged-dT modification was incorporated in a second 8×100 μl PCR containing 1X Q5 Reaction Buffer, 1X Q5 High GC Enhancer, 200 μM dNTPs, 250 nM PRA1_NoMod.F (Supplementary Table 1), 250 nM of an NPOM-caged-dT modified reverse primer that varied depending on target sequence (Supplementary Tables 1 and 2), 20 pM linear dsDNA template prepared as described above, and 0.02 U/μl Q5 DNA polymerase using the following thermal cycler protocol with a heated lid set to 105 °C: 98 °C for 30 s, [98 °C for 10 s, 65 °C for 30 s, 72 °C for 20 s] x 35 cycles, 72 °C for 2 min, hold at 12 °C. PCRs were then purified using a QIAquick PCR purification kit according to the manufacturer’s protocol and translesion DNA synthesis was performed by incubating three 100 μl reactions that contained 1X ThermoPol Buffer, 200 μM dNTPs, 0.02 U/μl *Sulfolobus* DNA polymerase IV, 0.02 U/μl Vent (exo-) DNA polymerase, and the purified PCR products at 55 °C for one hour. The resulting dsDNA templates were purified a second time using a QIAquick PCR purification kit according to the manufacturer’s protocol, eluted into 25 μl of 10 mM Tris (pH 8.0) per translesion synthesis reaction and quantified using the Qubit dsDNA Broad Range Assay kit with a Qubit 4 Fluorometer.

When preparing DNA templates by PCR, lambda exonuclease treatment, and primer extension, lambda exonuclease treatment was performed as 50 μl reactions containing 1X Lambda Exonuclease Reaction Buffer, up to 600 nM linear dsDNA template prepared as described above, and 0.1 U/μl lambda exonuclease. Reactions were incubated at 37 °C for 30 minutes, stopped by adding 1 μl of 500 mM EDTA (pH 8.5), and incubated at 75 °C for 10 min to heat inactivate lambda exonuclease. The sample volume was raised to 150 μl by adding 100 μl of 10 mM Tris (pH 8.0), mixed with an equal volume (150 μl) of phenol:chloroform:isoamyl alcohol (25:24:1) by vortexing, centrifuged at 18,000 x g and 4 °C for 5 min, and the aqueous supernatant was collected into a new tube. The samples were then ethanol precipitated by adding 0.1 volumes (15 μl) of 3M sodium acetate (pH 5.5), 3 volumes (450 μl) of 100% ethanol, and 1.5 μl of Glycoblue coprecipitant, and chilling at -20 °C overnight or -70 °C for 30-60 minutes. The samples were centrifuged at 18,000 x g and 4 °C for 30 min, the supernatant was discarded, and the samples were washed by adding 1 ml of 70% ethanol and inverting the tube gently. The samples were centrifuged at 18,000 x g and 4 °C for 5 min and the supernatant was discarded. The samples were briefly spun in a mini centrifuge, and residual liquid was discarded. Each pellet was then resuspended in 50 μl of 10 mM Tris (pH 8.0). The NPOM-caged-dT modification was then incorporated by converting the purified ssDNA into dsDNA in three 100 μl primer extension reactions containing 1X ThermoPol Buffer, 200 μM dNTPs, 300 μM NPOM-caged-dT modified primer (Supplementary Tables 1 and 2), ssDNA that was generated and purified as described above, and 0.02 U/μl Vent (exo-) DNA polymerase using the thermal cycler program: 95 °C for 3 min, 65 °C for 10 min, 72 °C for 10 min, hold at 12°C. 0.5 μl of thermolabile exonuclease I was added to each sample and the samples were incubated at 37 °C for 4 minutes and placed on ice. Thermolabile exonuclease I was then heat-inactivated by incubating the samples on a thermal cycler block that had been pre-heated to 80 °C for 1 minute. The primer extension reactions were purified using a QIAquick PCR purification kit according to the manufacturer’s protocol, eluted into 25 μl of 10 mM Tris (pH 8.0) per primer extension reaction and quantified using the Qubit dsDNA Broad Range Assay kit with a Qubit 4 Fluorometer. The sequences of all DNA templates are provided in Supplementary Table 3.

### Denaturing PAGE analysis of reversible transcription roadblocking

Single-round *in vitro* transcription was performed as described below for TECprobe-LM except that the volume of the transcription reaction was 87.5 μl and all wash and resuspension volumes were scaled accordingly. 25 μl aliquots of the transcription reaction were removed, transferred to 75 μl of TRIzol LS, and vortexed after the initial phase of transcription (pre-wash sample), after NTPs were depleted by washing (pre-chase sample), and after RNAP was chased to the terminal biotin-streptavidin roadblock (post-chase sample). 20 μl of chloroform was added to each sample and the samples were mixed by vortexing and inversion. The samples were centrifuged at 18,000 x g for 5 min and the aqueous phase was collected, mixed with 1.2 μl of GlycoBlue Coprecipitant and 50 μl of ice-cold isopropanol, incubated at room temperature for 15 min, and centrifuged at 18,000 x g for 15 min. The supernatant was discarded and the resulting pellet was washed with 200 μl of ice cold 70% ethanol by gently inverting the tube. The samples were centrifuged at 18,000 x g for 2 min and the supernatant was discarded. The samples were briefly spun in a mini centrifuge to pull down residual liquid, and the residual liquid was discarded. The samples were resuspended in 25 μl of 1X Turbo DNase Buffer, mixed with 0.75 μl of Turbo DNase, and incubated at 37 °C for 15 minutes. Each sample was mixed with 75 μl of TRIzol LS and purified by TRIzol extraction and isopropanol precipitation as described above. The resulting pellets were resuspended in 15 μl of Formamide Loading Dye (90% v/v deionized formamide, 1X Transcription Buffer (defined below), 0.05% w/v bromophenol blue), denatured by incubating at 95 °C for 5 min and run on an 8% polyacrylamide gel prepared using the SequaGel 19:1 Denaturing Gel system (National Diagnostics) for a Mini-PROTEAN Tetra Vertical Electrophoresis Cell. As described previously, denaturing conditions were achieved by filling the outer buffer chamber so that buffer covered only ∼1 cm of the gel plates, pre-running the gel at 480 V for 30 min, and running the gel at 480 V for ∼10 min^39^. Gels were stained with 1X SYBR Gold Nucleic Acid Stain in 1X TBE for 10 min and scanned on a Typhoon RGB Biomolecular Imager.

### TECprobe-LM

All steps of the TECprobe-LM procedure were performed under 592 nm amber light until the samples were irradiated with 365 nm UV light. For each set of TECprobe-LM samples (including pre-wash/pre-UV, pre-chase, and post-chase samples) a 165 μl single-round *in vitro* transcription reaction containing 1X Transcription Buffer (20 mM Tris-HCl (pH 8.0), 50 mM KCl, 1 mM DTT, 0.1 mM EDTA (pH 8.0)), 0.1 mg/ml BSA, 0.05% Tween-20, 10 nM DNA template, and 0.024 U/μl *E. coli* RNAP holoenzyme was prepared on ice and incubated at 37 °C for 15 min to form open promoter complexes. A 150 μl aliquot of 2 μg/μl streptavidin-coated magnetic beads that were equilibrated exactly as described previously for the TECprobe-VL procedure^20^ were placed on a magnetic stand and the supernatant was removed. The beads were then resuspended using the transcription reaction and incubated at room-temperature with end-over-end rotation at 15 rpm for 15 minutes to immobilize the template DNA. The sample was then briefly spun in a mini centrifuge and placed onto a magnetic stand for 1 minute to collect the beads on the tube wall. The supernatant was removed and the beads were resuspended in 660 μl of Wash Buffer 1 (1X Transcription Buffer, 0.05% Tween-20, 0.1 mg/ml BSA), returned to the magnetic stand, and the supernatant was removed. The beads were then washed using 660 μl of Wash Buffer 1 a second time. The beads were resuspended in 148.5 μl of Reaction Buffer 1 and incubated at 37 °C for 2 min before transcription was initiated by adding 16.5 μl of 10X Start Solution (100 mM MgCl_2_ and 0.1 mg/ml rifampicin). Upon addition of 10X start solution, the transcription reaction contained 1X Transcription Buffer, 0.1 mg/ml BSA, 0.05% Tween-20, 50 μM or 100 μM NTPs, 10 mM MgCl_2_, and 10 μg/μl rifampicin. In ZTP riboswitch experiments, the transcription reaction also contained 1 mM ZMP and 2% DMSO (1 mM ZMP samples) or 2% DMSO (0 mM ZMP samples). In fluoride riboswitch experiments, the transcription reaction also contained 10 mM NaF when fluoride was present. The transcription reaction was incubated at 37 °C for 2 min to allow RNAP to transcribe to the NPOM-caged-dT modification.

In all experiments except that of Figure 3, the pre-wash sample was then chemically probed by transferring 25 μl of the transcription reaction to a tube containing 2.78 μl of 400 mM benzoyl cyanide^41,42^ (BzCN, modified channel) and transferring an additional 25 μl to a tube containing 2.78 μl of 100% DMSO (untreated channel). Each channel of the pre-wash sample was then mixed with 75 μl of TRIzol LS and vortexed thoroughly. The remaining transcription reaction was placed on a magnetic stand, the supernatant was discarded, and the beads were resuspended in 840 μl of Wash Buffer 2. For all samples, Wash Buffer 2 contained 1X Transcription Buffer, 0.1 mg/ml BSA, 0.05% Tween-20, 10 mM MgCl_2_, and 10 μg/ml rifampicin. For ZTP riboswitch experiments performed in the presence of ZMP, Wash Buffer 2 also contained 0.1 mM ZMP and 0.2% DMSO. For ZTP riboswitch experiments performed in the absence of ZMP, Wash Buffer 2 also contained 0.2% DMSO. For fluoride riboswitch experiments performed in the presence of fluoride, Wash Buffer 2 also contained 1 mM NaF. The beads were transferred to a new tube, placed on a magnetic stand, and the supernatant was removed. The beads were resuspended in 840 μl of Wash Buffer 2 a second time, transferred to a new tube, placed on a magnetic stand, and the supernatant was removed. The beads were then resuspended in 840 μl of Wash Buffer 2 a third time, transferred to a new tube, incubated on an end-over-end rotator at room temperature for 5 min, briefly spun down in a mini centrifuge, placed on a magnetic stand, and the supernatant was removed. The beads were then resuspended in 113.7 μl of Reaction Buffer 2 which, upon completing the reaction by adding NTPs in a later step, contained 1X Transcription Buffer, 0.1 mg/ml BSA, 0.05% Tween-20, 10 mM MgCl_2_, 10 μg/ml rifampicin. For ZTP riboswitch experiments, Reaction Buffer 2 also contained 1 mM ZMP and 2% DMSO (1 mM ZMP samples) or 2% DMSO (0 mM ZMP samples). For fluoride riboswitch experiments, Reaction Buffer 2 also contained 10 mM NaF when fluoride was present.

In the experiment shown in Figure 3 in which three-point chemical probing was performed, samples were washed immediately after RNAP had transcribed to the NPOM-caged-dT roadblock as follows: The sample was placed on a magnetic stand and the supernatant was discarded. The beads were resuspended in 840 μl of Wash Buffer 2, transferred to a new tube, incubated on an end-over-end rotator at room temperature for 5 min, briefly spun down in a mini centrifuge, placed on a magnetic stand, and the supernatant was removed. The beads were then resuspended in 840 μl of Wash Buffer 2 and washed as described above a second time. For experiments performed in the presence of ZMP, Wash Buffer 2 was supplemented with 0.1 mM ZMP and 0.2% DMSO. For experiments performed in the absence of ZMP, Wash Buffer 2 was supplemented with 0.2%DMSO. The beads were then resuspended in 163.7 μl of Reaction Buffer 2 and incubated at 37 °C for 2 min. The pre-UV sample was then chemically probed as described above.

The sample was placed on a custom-built microcentrifuge tube irradiator^39^ and irradiated with 10 mW/cm^2^ 365 nm UV light for 3 minutes, with mixing after every minute. The sample was incubated at 37 °C for 2 min and the pre-chase sample was chemically probed and stopped with TRIzol LS as described above for the pre-wash sample. 1.3 μl of 5 mM NTPs were added to the remaining sample and the reaction was incubated at 37 °C for 2 min to allow RNAP to transcribe to the terminal biotin-streptavidin roadblock. The post-chase sample was then chemically probed and stopped with TRIzol LS as described above for the pre-wash sample.

The samples were then TRIzol extracted and converted to cDNA and indexed dsDNA Illumina libraries by ligating an adapter to the RNA 3’ end, performing solid-phase error-prone reverse transcription, and degrading the RNA using the TECprobe-VL protocol for sequencing library preparation, which was described in detail previously^20^.

### TECprobe-VL

The preparation of randomly biotinylated DNA templates, TECprobe-VL procedure, and analysis of TECprobe-VL data were performed exactly as described previously^20^ except that whole-dataset normalization was performed using process_TECprobeVL_profiles as described below.

### High-throughput DNA sequencing

Sequencing was performed by Novogene Co. on an Illumina HiSeq X Ten System using 2×150 PE reads with 10% PhiX spike-in. TECprobe-LM libraries were sequenced to a depth of ∼10 million PE reads. TECprobe-VL libraries were sequenced to a depth of ∼45 to ∼60 million PE reads.

### Sequencing read pre-processing, alignment, and SHAPE-MaP reactivity calculation

All custom software are freely available at https://github.com/e-strobel-lab/TECtools/releases/tag/v1.2.0 or https://github.com/e-strobel-lab/TECprobe_visualization/releases/tag/v1.0.0.

Because the TECprobe-LM sequencing library preparation is identical to that of TECprobe-VL, sequencing read pre-processing and alignment were performed exactly as described previously^20^. Briefly, 3’ end targets and intermediate transcript targets were generated by running `cotrans_preprocessor` in `MAKE_3pEND_TARGETS` mode. Sequencing reads were then processed by running `cotrans_preprocessor` in `PROCESS_MULTI` mode, which manages adapter trimming using fastp^43^ and demultiplexes sequencing reads by 3’ end identity and channel (modified or untreated). A shell script to run ShapeMapper2^44^ for every intermediate transcript was generated by running `cotrans_preprocessor` in `MAKE_RUN_SCRIPT` mode. Sequencing read alignment and reactivity calculation was then performed by ShapeMapper2.

During ShapeMapper2^44^ analysis, TECprobe-LM and TECprobe-VL reactivity profiles are normalized individually for each transcript because sequencing reads are demultiplexed by 3’ end identity prior to analysis to avoid multi-mapping. Reactivity profiles for each transcript were therefore normalized together using `process_TECprobeVL_profiles`, which supersedes the `compile_SM2_output` script. The minimum depth required for a nucleotide to be used when calculating the normalization factor was set to 50,000, the maximum background mutation rate was 0.05, nucleotides masked by lowercase were excluded, and the left-most nucleotide was excluded. Normalization was performed exactly as done by ShapeMapper2^44,45^: The reactivity of each nucleotide was divided by the mean reactivity of the top 10% of reactivity values after excluding reactivity values above the largest value out of: i) 1.5x the interquartile range or ii) the 90^th^ or 95^th^ percentile, depending on whether most high-quality reactivity values originated from transcripts in which the target RNA length was >100 or <100, respectively. `process_TECprobeVL_profiles` generates a directory that contains i) a record of input and output file names, ii) sub-directories that contain ShapeMapper2 reactivity profile files for every transcript length in which normalized reactivity values were computed using the whole data set, and iii) a csv file in which data from the ShapeMapper2 profiles for each transcript length are assembled into matrices, which is formatted identically to the output of `compile_sm2_output`. For TECprobe-LM data, each sample was processed by `process_TECprobeVL_profiles` individually. For the *E. coli* SRP RNA, three replicates were provided to `process_TECprobeVL_profiles` together, which merges the data prior to normalization factor calculation. The *E. coli* SRP RNA reactivity heatmap was generated using `generate_cotrans_heatmap`. In the *C*. *beijerinckii pfl* ZTP riboswitch datasets, the reactivity of C98 was masked as 0 because its background mutation rate was either close to or above the maximum background mutation rate of 0.05.

## Data availability

The raw sequencing data generated in this study have been deposited in the Sequencing Read Archive (https://www.ncbi.nlm.nih.gov/sra) with the BioProject accession code PRJNA992462. Individual BioSample accession codes are available in Supplementary Table 4. Fully processed reactivity data have been deposited in the RNA Mapping Database^46^ (https://rmdb.stanford.edu/). Individual accession codes for each data set are available in Supplementary Table 5. ShapeMapper2 output files that were re-normalized using process_TECprobeVL_profiles have been deposited in Zenodo (DOI: 10.5281/zenodo.13871546). TIFF images of all gels generated in this study have been deposited in Zenodo (DOI: 10.5281/zenodo.13871636).

## Code availability

TECtools can be accessed at https://github.com/e-strobel-lab/TECtools/releases/tag/v1.2.0.

## Acknowledgements

This work was supported by the National Institute of General Medical Sciences of the National Institutes of Health under Award Number R35GM147137 (to E.J.S) and by start-up funding from the University at Buffalo (to E.J.S). The content is solely the responsibility of the authors and does not necessarily represent the official views of the National Institutes of Health.

## Author Contributions

E.J.S., conceptualization; C.E.S. and E.J.S., methodology; C.E.S. and S.L.K., investigation; C.E.S. and E.J.S, Validation; C.E.S., S.L.K., and E.J.S, Formal Analysis; E.J.S, Software; E.J.S, writing – original draft; C.E.S., S.L.K., and E.J.S, writing – review & editing; E.J.S., supervision; E.J.S., funding acquisition.

---

**Supplementary Figure 1.**
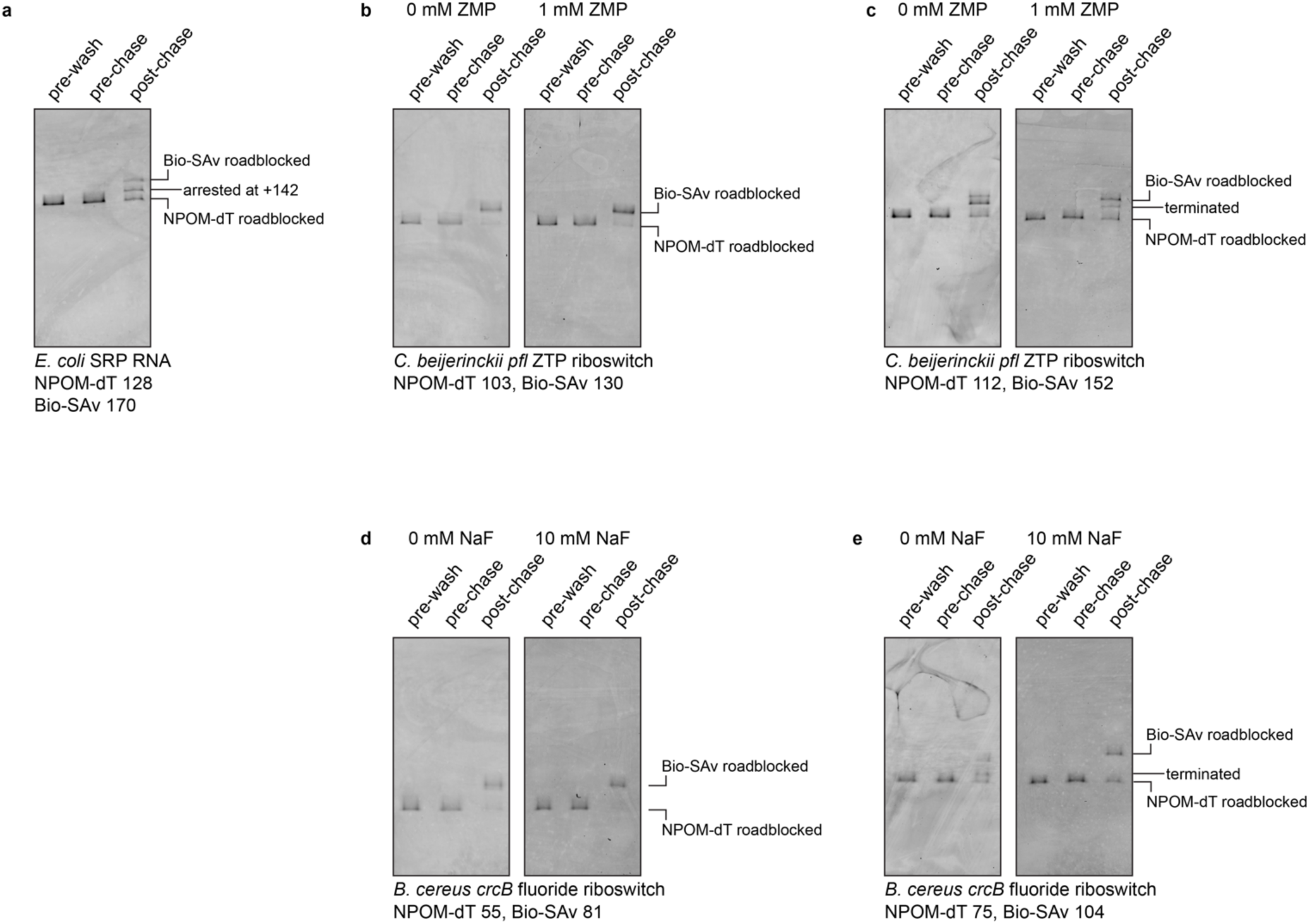
Analysis of reversible transcription roadblocking by denaturing gel electrophoresis. Denaturing PAGE analysis of transcripts generated during the TECprobe-LM transcription reactions for the (**a**) *E. coli* SRP RNA, (**b**, **c**) *C. beijerinckii pfl* ZTP riboswitch, and (**d** and **e**) *B. cereus crcB* fluoride riboswitch. Bio-SAv, biotin-streptavidin.

**Supplementary Figure 2.**
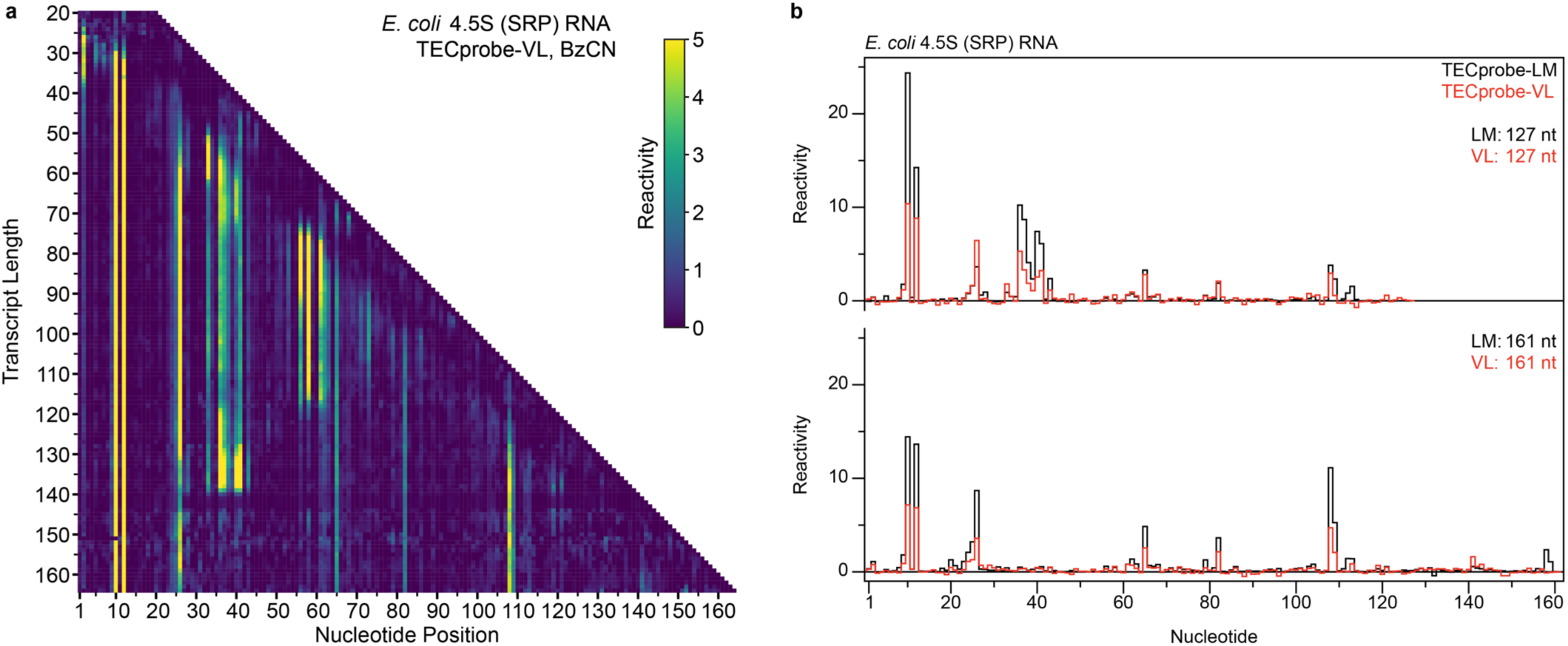
Comparison of TECprobe-LM and TECprobe-VL reactivity profiles for the *E. coli* SRP RNA. (**a**) TECprobe-VL reactivity matrix for the *E*. *coli* SRP RNA. (**b**) Comparison of TECprobe-LM and TECprobe-VL reactivity profiles for the 127 and 161 nt transcripts of the *E. coli* SRP RNA. TECprobe-LM data are the average of n=2 replicates; n=3 TECprobe-VL replicates were concatenated and processed together. SRP, signal recognition particle; BzCN, benzoyl cyanide.

**Supplementary Figure 3.**
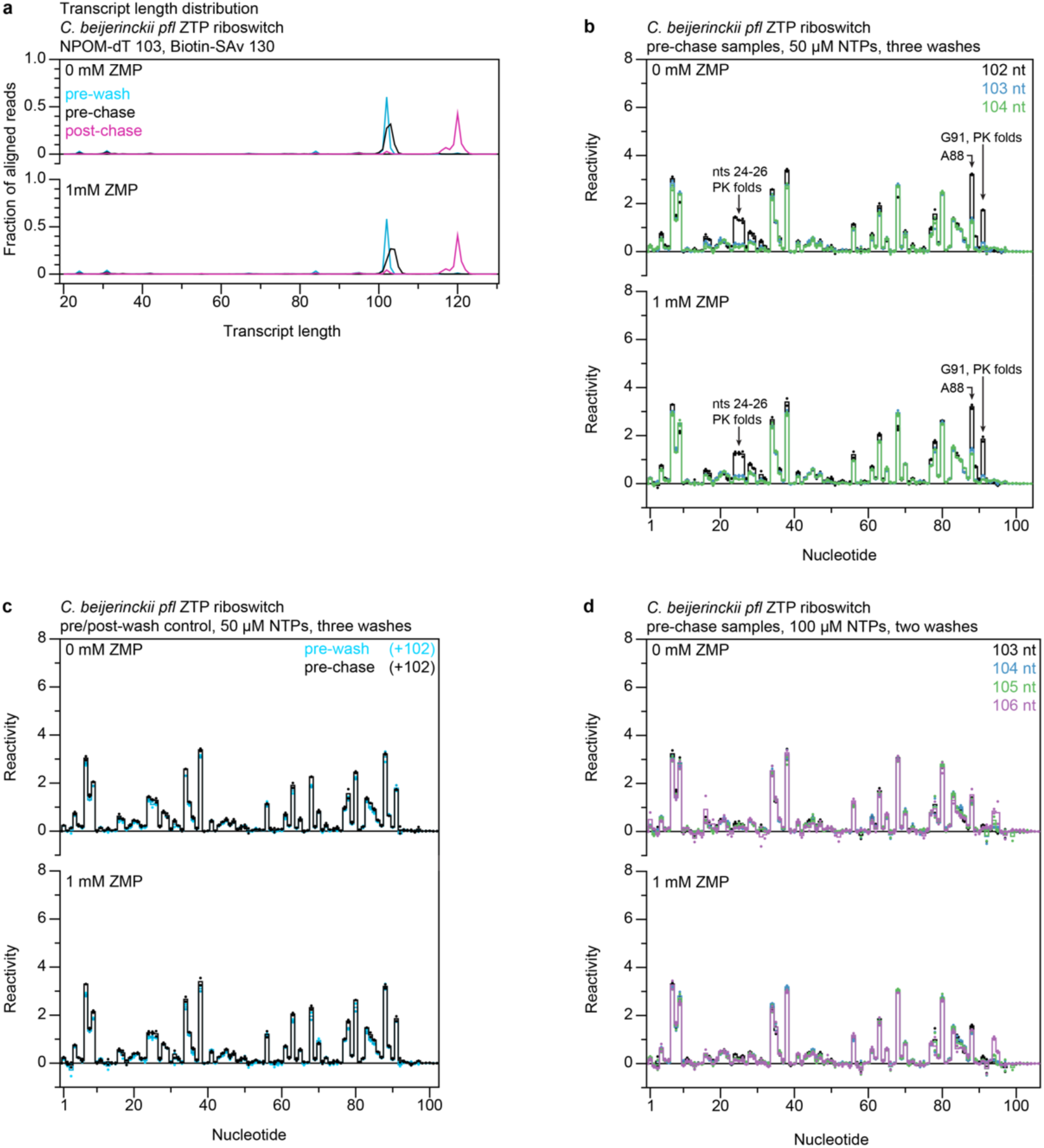
Additional analyses of *C. beijerinckii pfl* ZTP riboswitch aptamer TECprobe-LM data. (**a**) Transcript length distribution for the pre-wash, pre-chase, and post-chase samples when transcription was performed with 50 μM NTPs and three washes. Traces are the average of n=2 replicates. (**b**) Pre-chase reactivity profiles for the 102, 103, and 104 nt transcripts from the TECprobe-LM experiments that were performed with 50 μM NTPs and three washes. (**c**), Comparison of pre-wash and pre-chase reactivity profiles for the 102 nt transcript when TECprobe-LM was performed using 50 μM NTPs and three washes. (**d**) Pre-chase reactivity profiles for 103, 104, 105, and 106 nt transcripts from the TECprobe-LM experiments that were performed with 100 μM NTPs and limited washing. In b-d, solid lines are the average of n=2 replicates and reactivity values for individual replicates are shown as points. SAv, streptavidin; PK, pseudoknot.

**Supplementary Figure 4.**
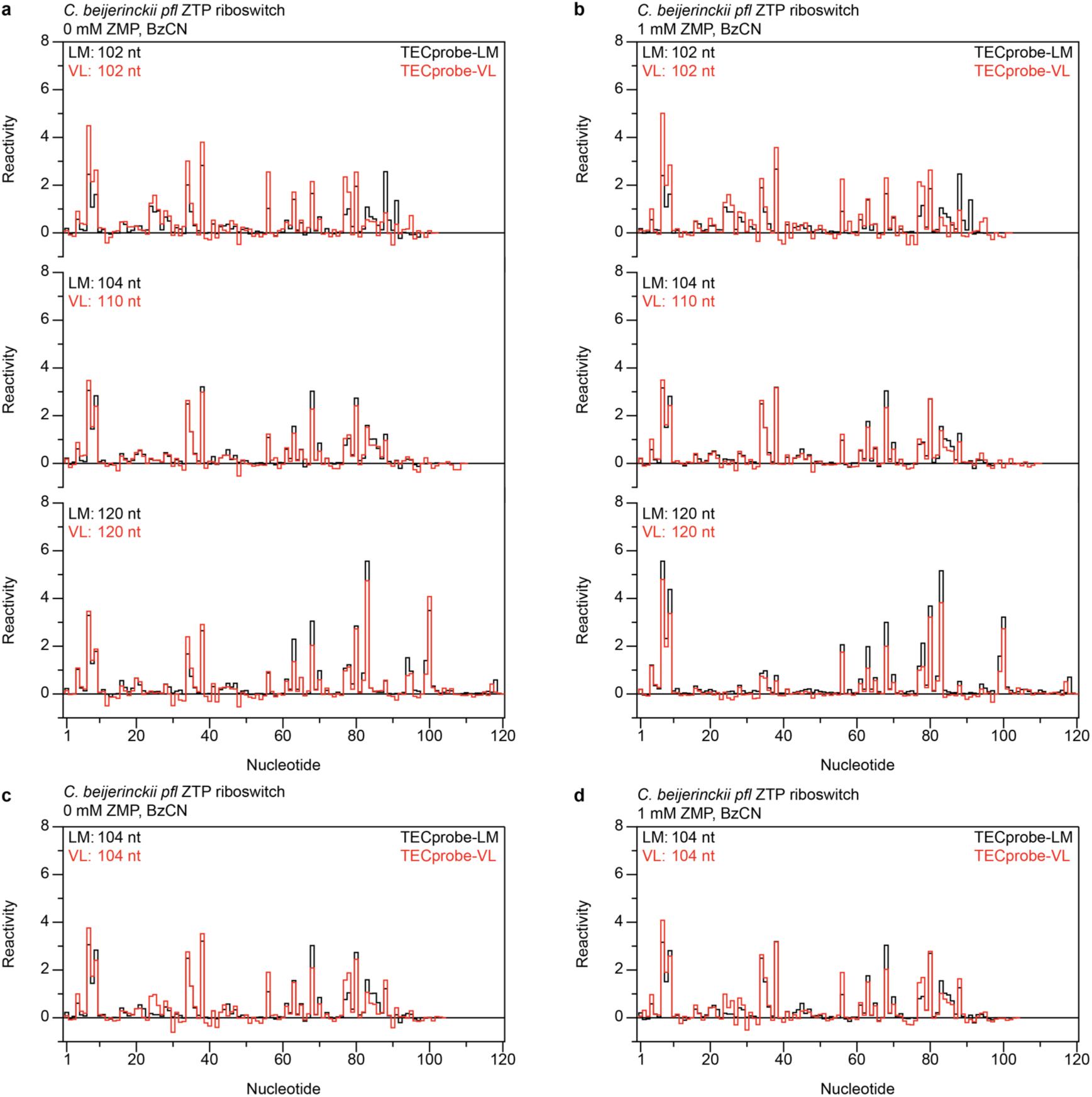
Comparison of TECprobe-LM and TECprobe-VL reactivity profiles for *C. beijerinckii pfl* ZTP riboswitch aptamer folding intermediates. (**a, b**) Comparison of TECprobe-LM and TECprobe-VL reactivity profiles for the *C. beijerinckii pfl* ZTP riboswitch pseudoknot formation and ligand binding folding transitions. The TECprobe-LM profile of the 104 nt transcript was compared to the TECprobe-VL profile of the 110 nt transcript because the pseudoknot does not completely fold until 110 nt in the TECprobe-VL data set, presumably due to biotin-streptavidin induced backtracking. The TECprobe-LM and TECprobe-VL profiles for the 104 nt transcript are compared in panels (**c**) and (**d**). TECprobe-LM data are the average of n=2 replicates. TECprobe-VL data are from Szyjka and Strobel, *Observation of coordinated RNA folding events by systematic cotranscriptional RNA structure probing, Nat Commun.* 2023 Nov 29; 14(1):7839. n=2 TECprobe-VL replicates were concatenated and processed together. BzCN, benzoyl cyanide.

**Supplementary Figure 5.**
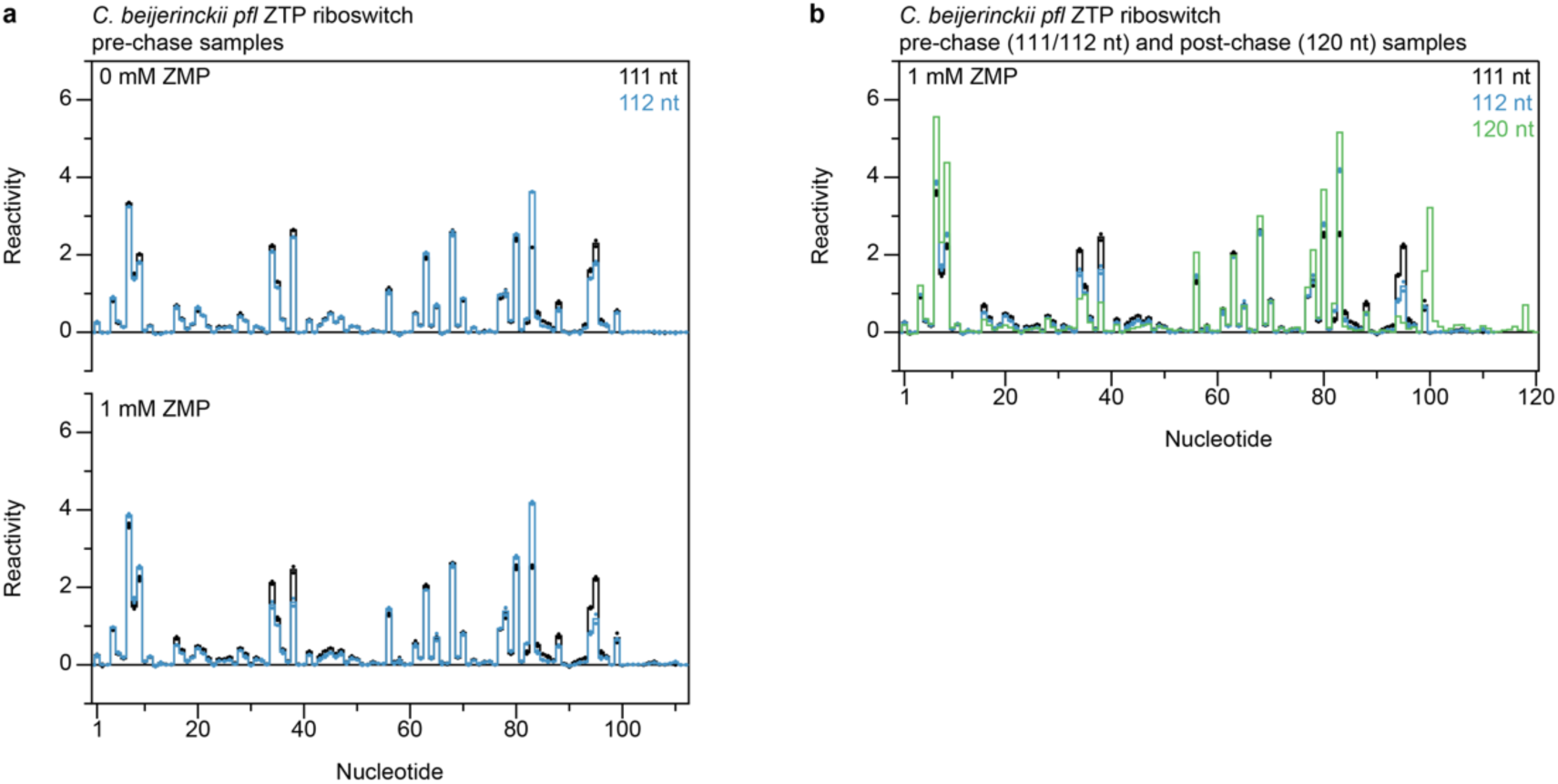
Additional analyses of *C*. *beijerinckii pfl* ZTP riboswitch expression platform folding TECprobe-LM data. (**a**) Comparison of pre-chase reactivity profiles for the 111 and 112 nt transcripts. (**b**) Comparison of reactivity profiles for the 111 and 112 nt transcripts from the 1 mM ZMP *pfl* ZTP riboswitch expression platform folding dataset with the 120 nt transcript from the 1 mM ZMP *C*. *beijerinckii pfl* ZTP riboswitch aptamer folding dataset.

**Supplementary Figure 6.**
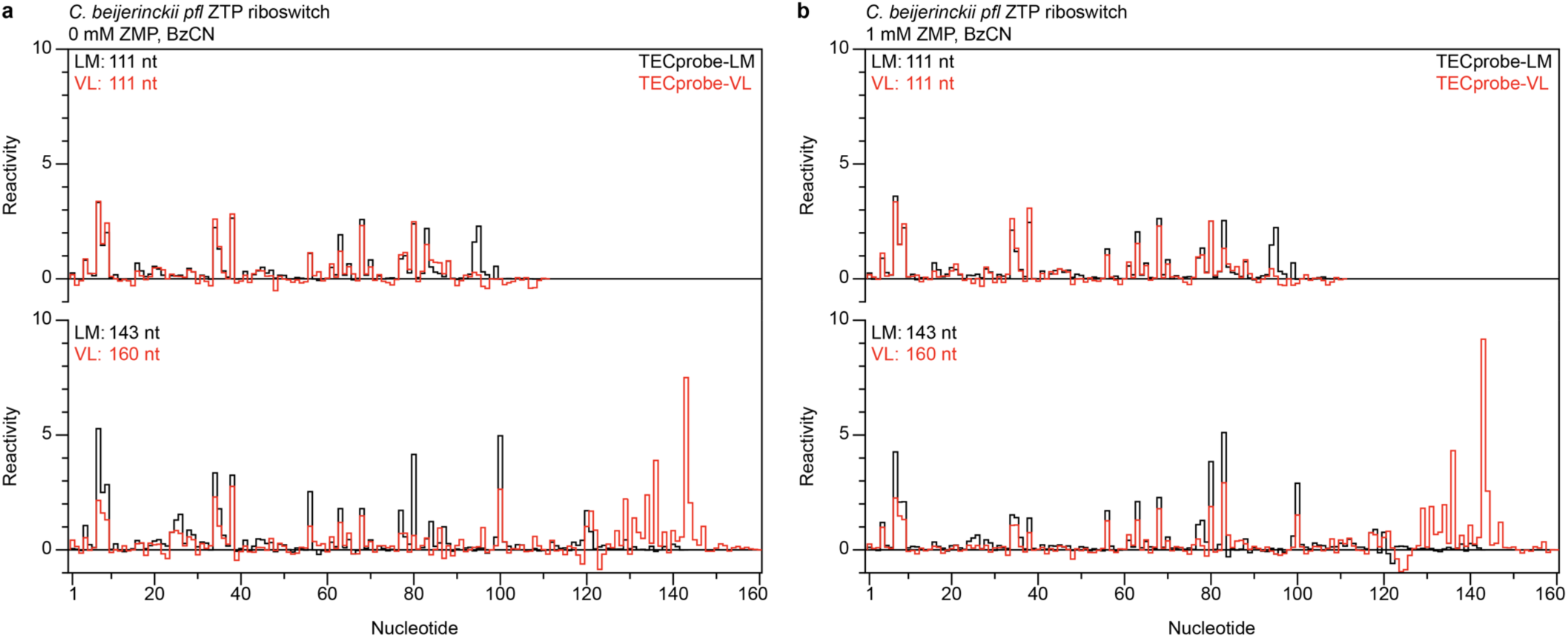
Comparison of TECprobe-LM and TECprobe-VL reactivity profiles for *C. beijerinckii pfl* ZTP riboswitch expression platform folding intermediates. Comparison of TECprobe-LM TECprobe-VL reactivity profiles for the *C. beijerinckii pfl* ZTP riboswitch aptamer-to-full length riboswitch folding transition with (**a**) 0 mM and (**b**) 1 mM ZMP. The TECprobe-LM profile of the 143 nt transcript was compared to the TECprobe-VL profile of the 160 nt transcript because the 143 nt transcript was not enriched in the TECprobe-VL sequencing library TECprobe-LM data are the average of n=2 replicates. TECprobe-VL data are from Szyjka and Strobel, *Observation of coordinated RNA folding events by systematic cotranscriptional RNA structure probing, Nat Commun.* 2023 Nov 29; 14(1):7839. n=2 TECprobe-VL replicates were concatenated and processed together. BzCN, benzoyl cyanide.

**Supplementary Figure 7.**
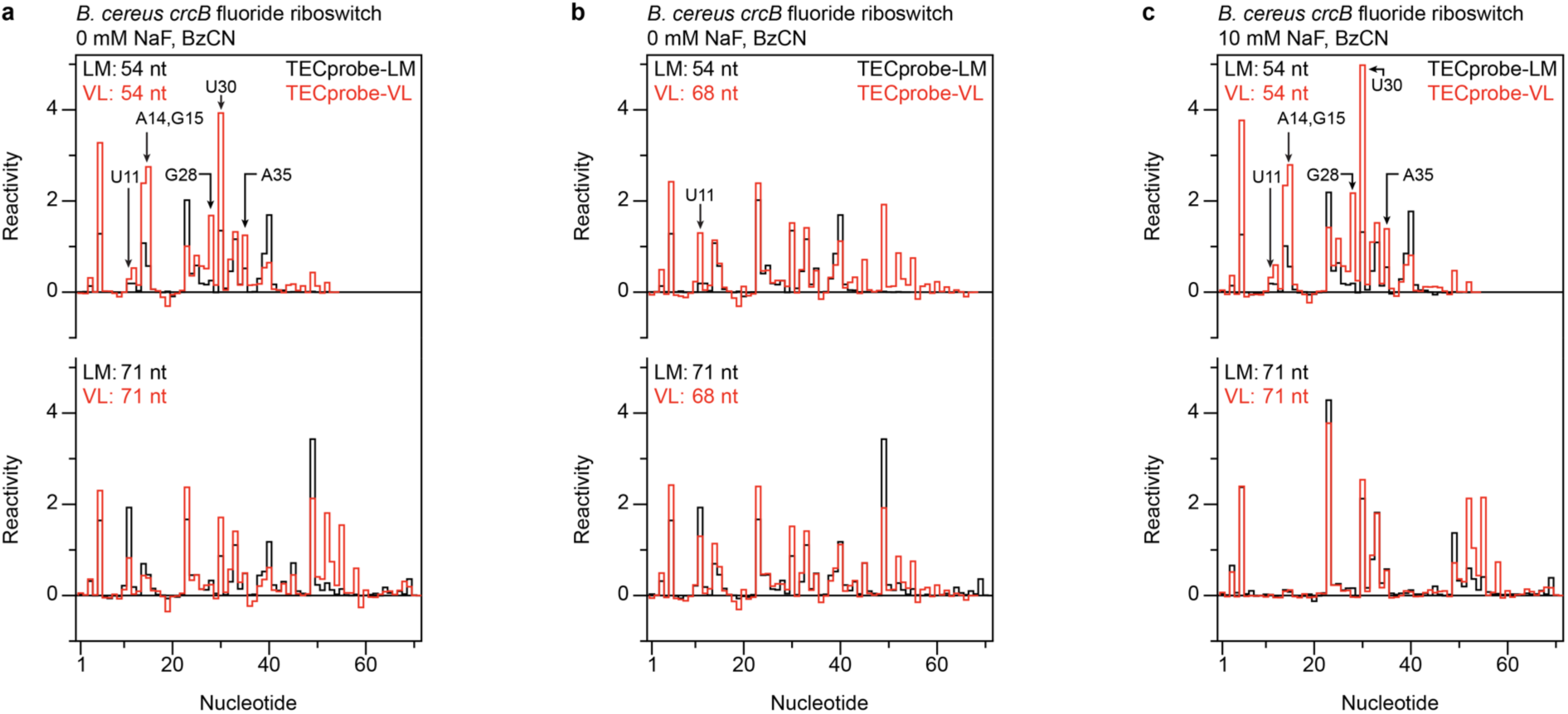
Comparison of TECprobe-LM and TECprobe-VL reactivity profiles for *B. cereus crcB* fluoride riboswitch aptamer folding intermediates. (**a**) Comparison of TECprobe-LM and TECprobe-VL reactivity profiles for the 54 and 71 nt transcripts of the *B. cereus crcB* fluoride riboswitch with 0 mM NaF. (**b**) Comparison of the 0mM NaF TECprobe-LM reactivity profiles for the 54 and 71 nt transcripts of the *B. cereus crcB* fluoride riboswitch with the 0 mM NaF TECprobe-VL reactivity profile for the 68 nt transcript. (**c**) Comparison of TECprobe-LM and TECprobe-VL reactivity profiles for the 54 and 71 nt transcripts of the *B. cereus crcB* fluoride riboswitch with 1 mM NaF.TECprobe-LM data are the average of n=2 replicates. TECprobe-VL data are from Szyjka and Strobel, *Observation of coordinated RNA folding events by systematic cotranscriptional RNA structure probing, Nat Commun.* 2023 Nov 29; 14(1):7839. n=2 TECprobe-VL replicates were concatenated and processed together. BzCN, benzoyl cyanide.

**Supplementary Figure 8.**
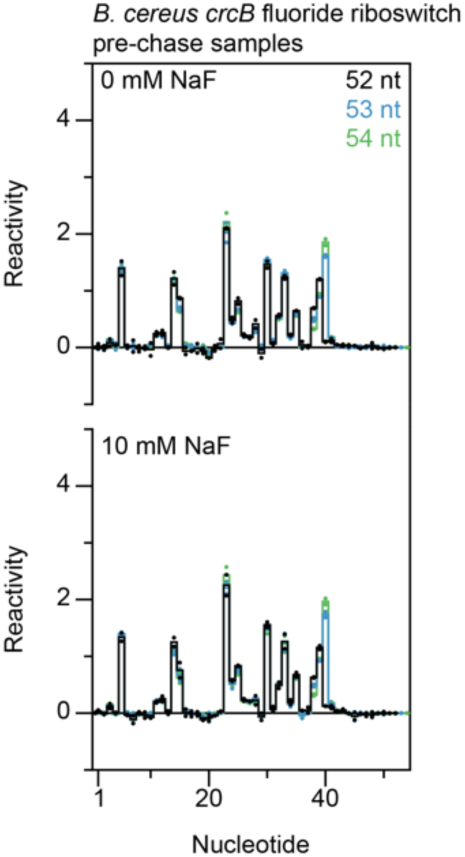
Additional analyses of *B. cereus crcB* fluoride riboswitch aptamer TECprobe-LM data. Comparison of pre-chase reactivity profiles for the 52, 53, and 54 nt transcripts. Solid lines are the average of n=2 replicates and reactivity values for individual replicates are shown as points.

**Supplementary Figure 9.**
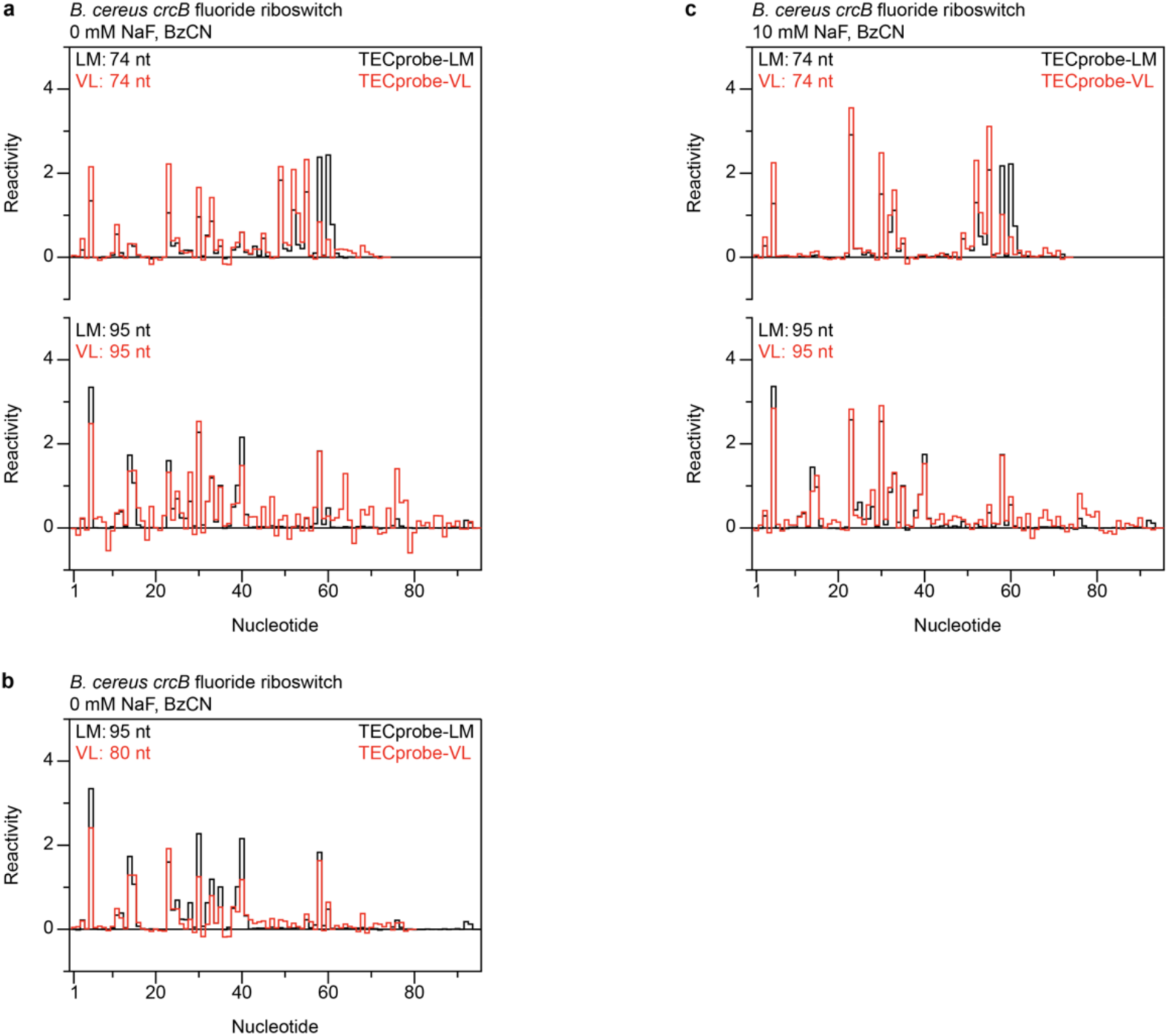
Comparison of TECprobe-LM and TECprobe-VL reactivity profiles for *B. cereus crcB* fluoride riboswitch expression platform folding intermediates. (**a**) Comparison of TECprobe-LM and TECprobe-VL reactivity profiles for the 74 and 95 nt transcripts of the *B. cereus crcB* fluoride riboswitch without NaF. (**b**) Comparison of the reactivity profiles for the 95 (TECprobe-LM) and 80 nt (TECprobe-VL) transcripts of the *B. cereus crcB* fluoride riboswitch without NaF. (**c**) Comparison of TECprobe-LM and TECprobe-VL reactivity profiles for the 74 and 95 nt transcripts of the *B. cereus crcB* fluoride riboswitch with 10 mM NaF. TECprobe-LM data are the average of n=2 replicates. TECprobe-VL data are from Szyjka and Strobel, *Observation of coordinated RNA folding events by systematic cotranscriptional RNA structure probing, Nat Commun.* 2023 Nov 29; 14(1):7839 n=2 TECprobe-VL replicates were concatenated and processed together. BzCN, benzoyl cyanide

**Supplementary Table 1.**
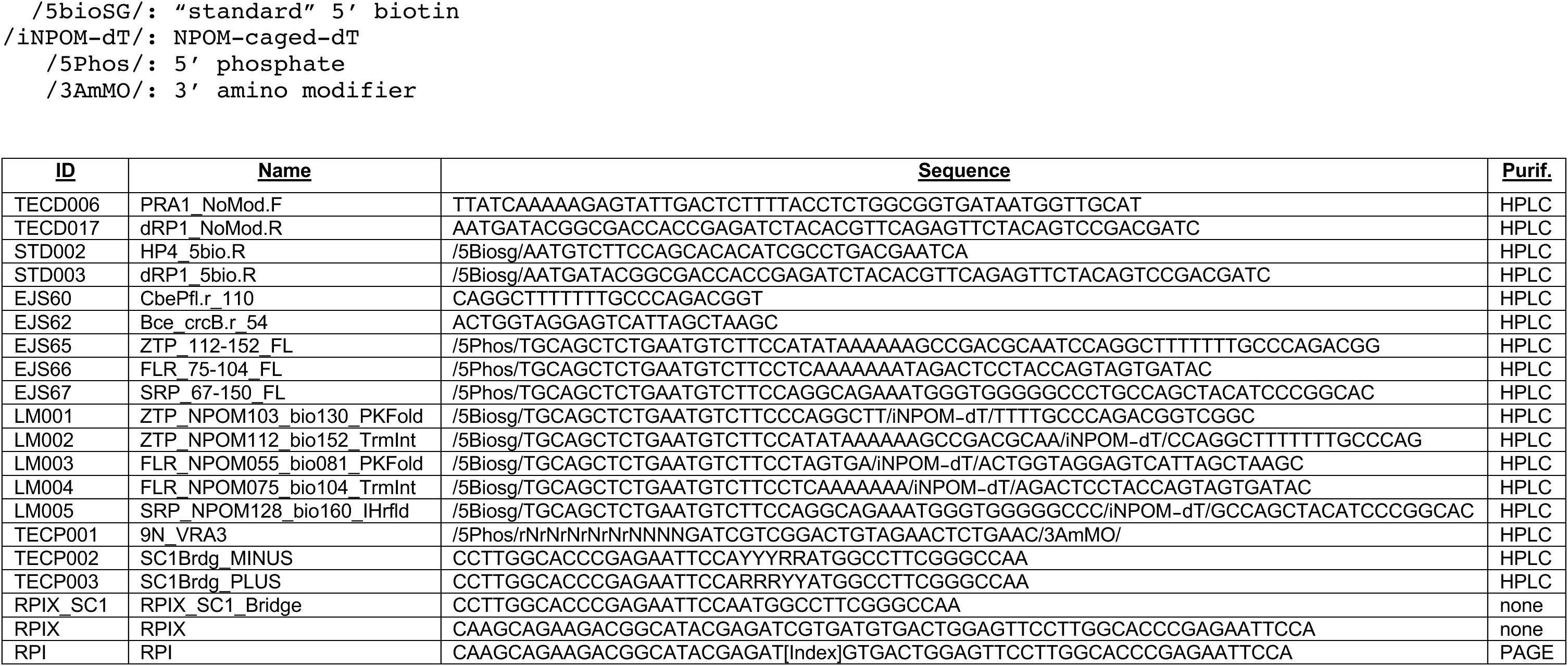
Oligonucleotides used in this study. The table below shows oligonucleotides used in this study. The modification codes presented are compatible with Integrated DNA Technology ordering.

**Supplementary Table 2.** DNA templates prepared for this study. The table below describes the DNA templates prepared for this study, including the primers and templates used, DNA modifications, whether translesion synthesis was performed, how the DNA template was purified, and the figures in which each DNA templates was used

| <u>ID</u> | <u>Fwd Primer</u> | <u>Rev Primer</u> | <u>Template</u> | <u>Modifications</u> | <u>Translesion Synthesis</u> | <u>Clean up</u> | <u>Used in Fig(s)</u> |
| --- | --- | --- | --- | --- | --- | --- | --- |
| 1 | TECD006 | EJS60 | pCES008 | None | N/A | Gel extracted | N/A |
| 2 | TECD006 | EJS65 | pCES008 | 5' phosphate | N/A | Gel extracted | N/A |
| 3 | TECD006 | EJS62 | pCES010 | None | N/A | Gel extracted | N/A |
| 4 | TECD006 | EJS66 | pCES010 | 5' phosphate | N/A | Gel extracted | N/A |
| 5 | TECD006 | EJS67 | pCES018 | 5' phosphate | N/A | Gel extracted | N/A |
| 6 | TECD006 | LM001 | Template 1 | Internal NPOM-caged-dT, 5' biotin | Yes | PCR purification columns | 3, S1, S3, S4, S5 |
| 7 | TECD006 | LM002 | Template 2 | Internal NPOM-caged-dT, 5' biotin | N/A | Phenol:chloroform extraction, PCR purification columns | 4, S1, S5, S6 |
| 8 | TECD006 | LM003 | Template 3 | Internal NPOM-caged-dT, 5' biotin | Yes | PCR purification columns | 5, S1, S7, S8 |
| 9 | TECD006 | LM004 | Template 4 | Internal NPOM-caged-dT, 5' biotin | N/A | Phenol:chloroform extraction, PCR purification columns | 6, S1, S9 |
| 10 | TECD006 | LM005 | Template 5 | Internal NPOM-caged-dT, 5' biotin | N/A | Phenol:chloroform extraction, PCR purification columns | 2, S2 |
| 11 | TECD006 | STD002 | pCES018 | 5' biotin | N/A | Gel extracted | N/A |
| 12 | TECD006 | STD002 | Template 11 | Internal biotin-11 nucleotides, 5' biotin | N/A | SPRI beads | S2 |

**Supplementary Table 3.**
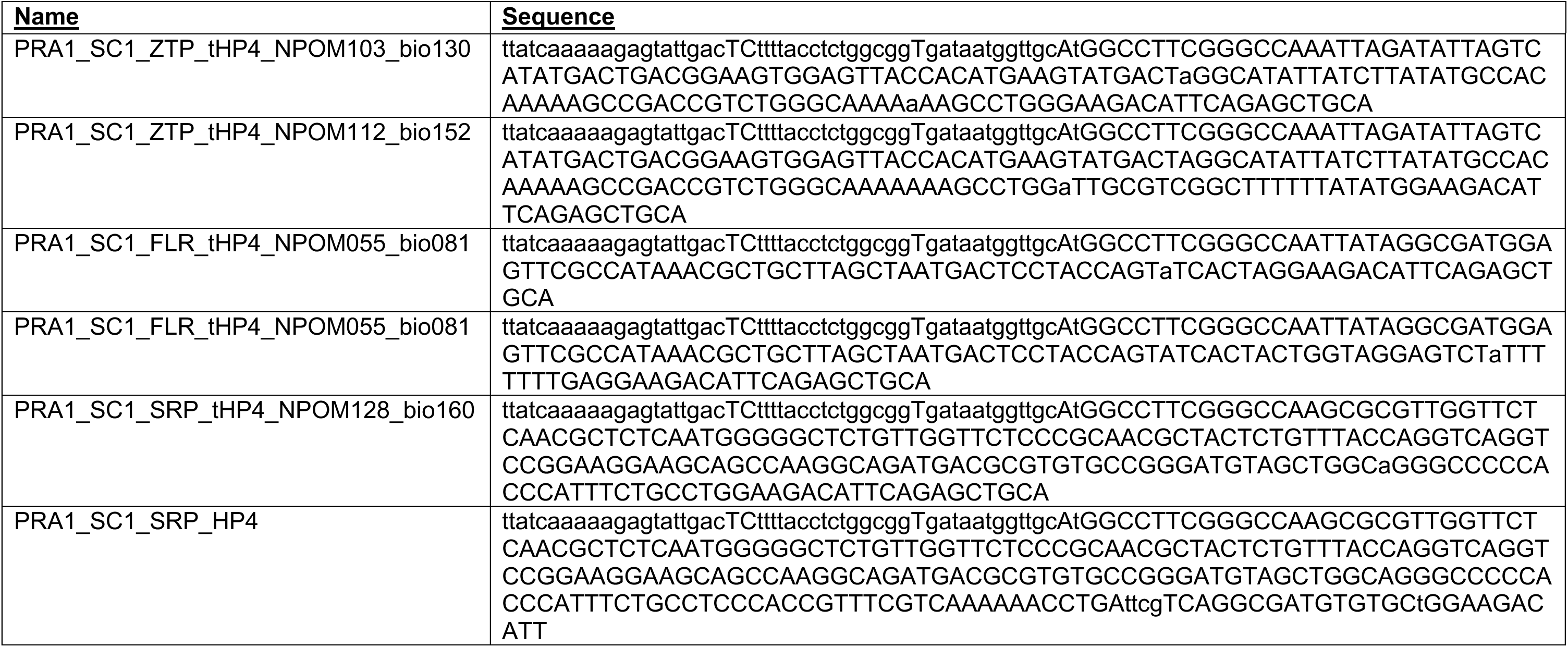
DNA template sequences. The table below contains the DNA sequences used in this study. The location of the NPOM-caged-dT stall site is indicated by a lowercase ‘a’ with in the uppercase target RNA sequence.

**Supplementary Table 4.**
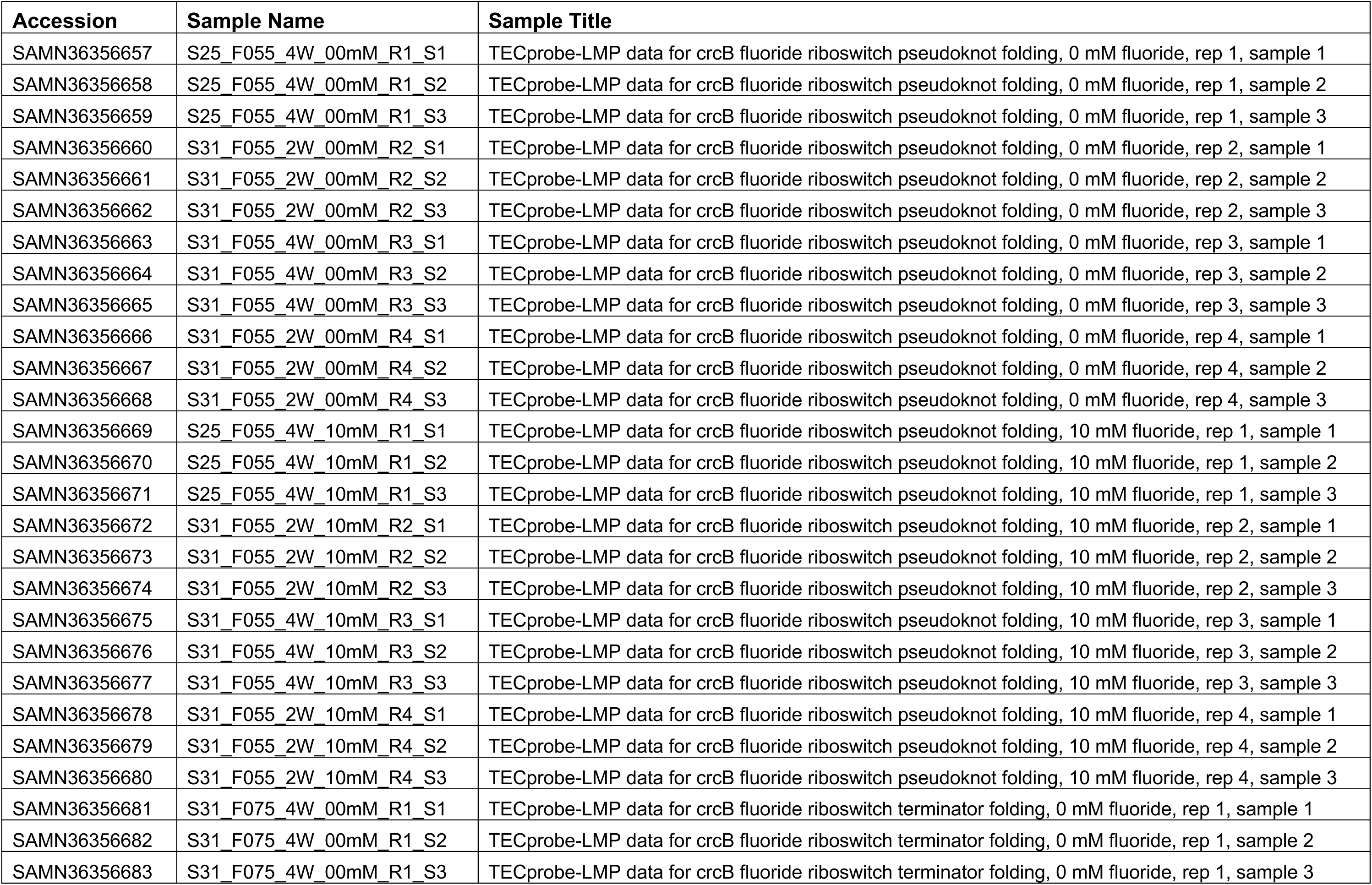
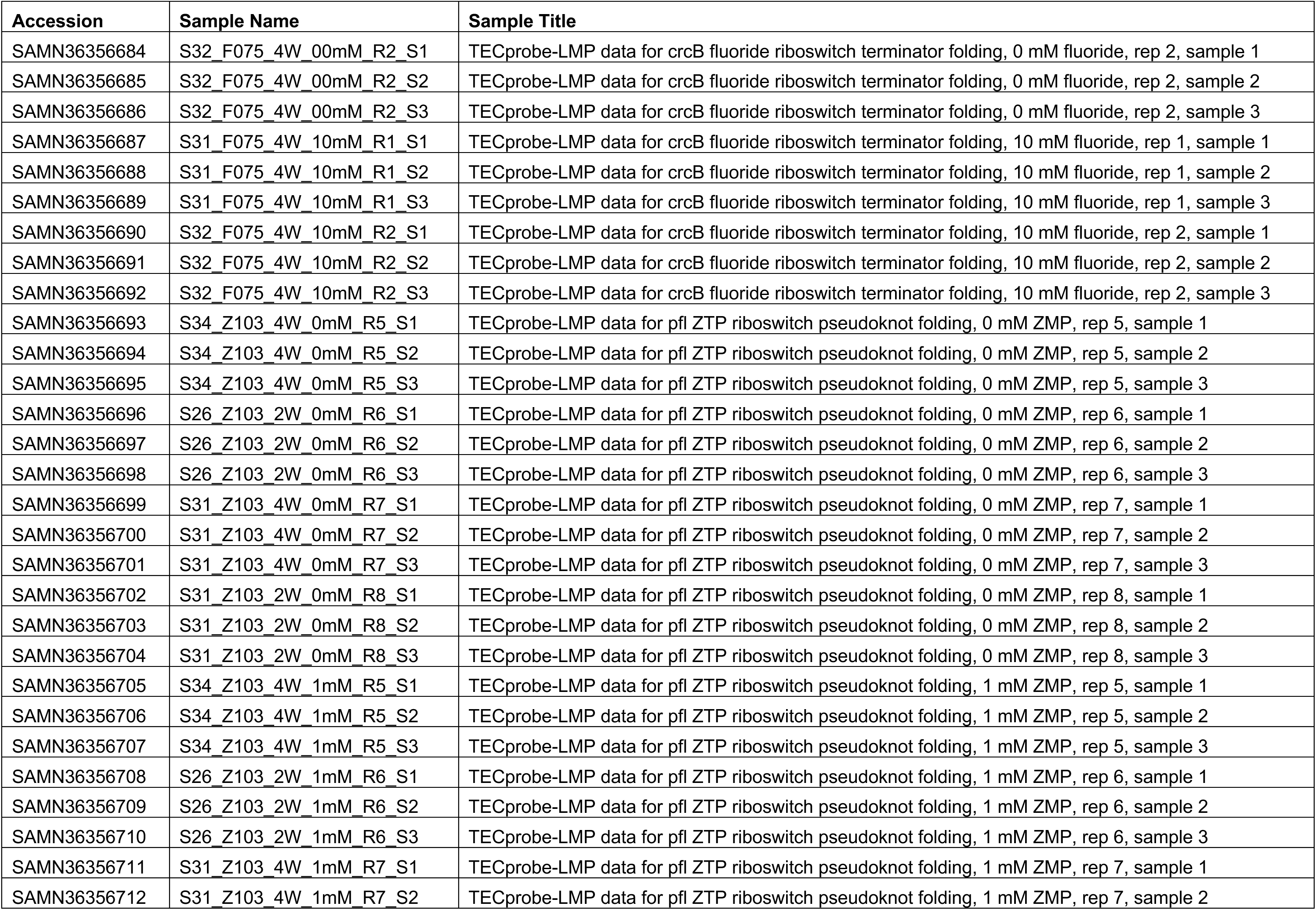
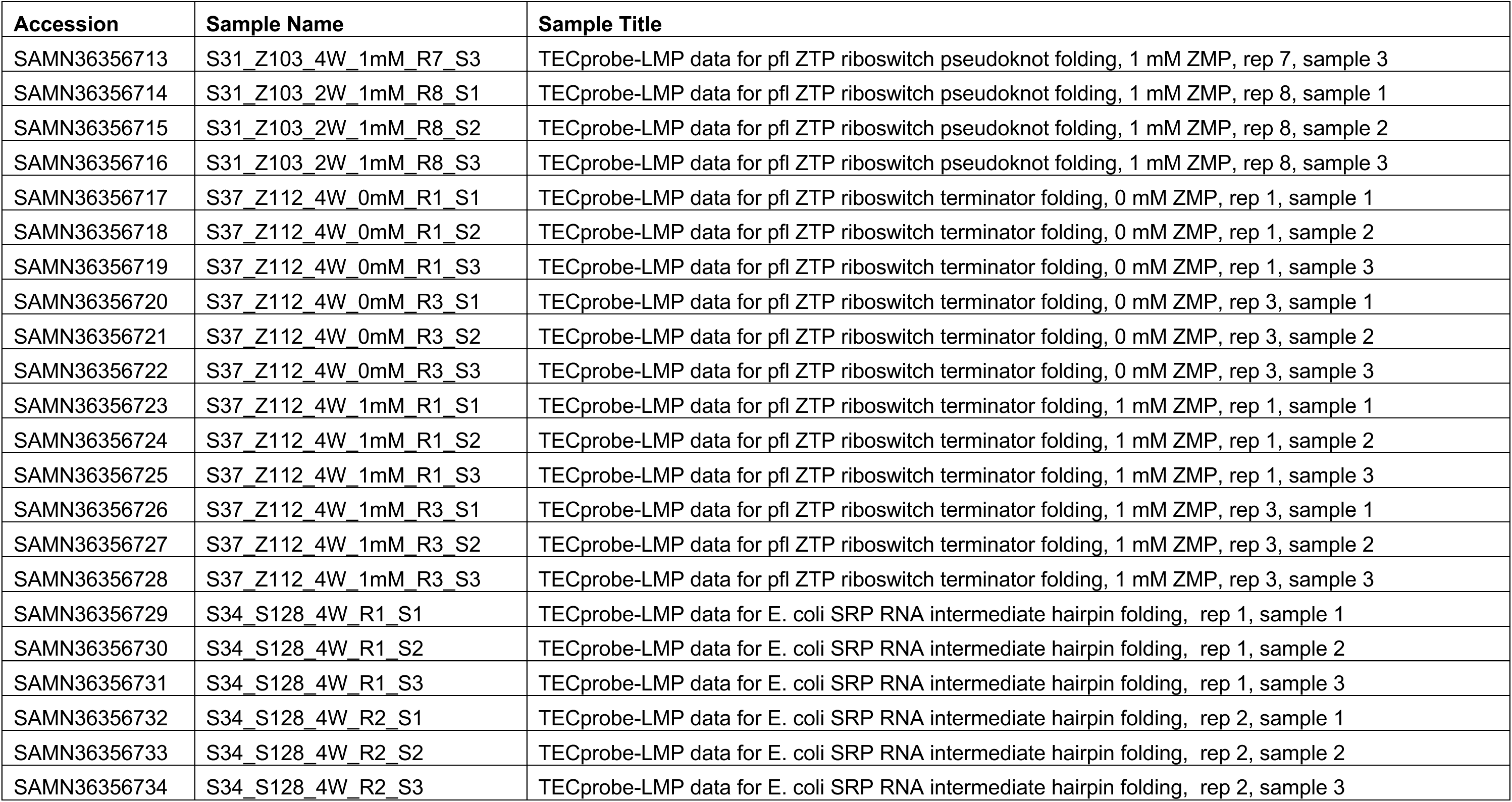
Sequencing Read Archive (SRA) deposition table. All primary sequencing data generated in this work are freely available from the Sequencing Read Archive (http://www.ncbi.nlm.nih.gov/sra), accessible via the BioProject accession number <u>PRJNA992462</u> or using the individual accession numbers below.

**Supplementary Table 5.**
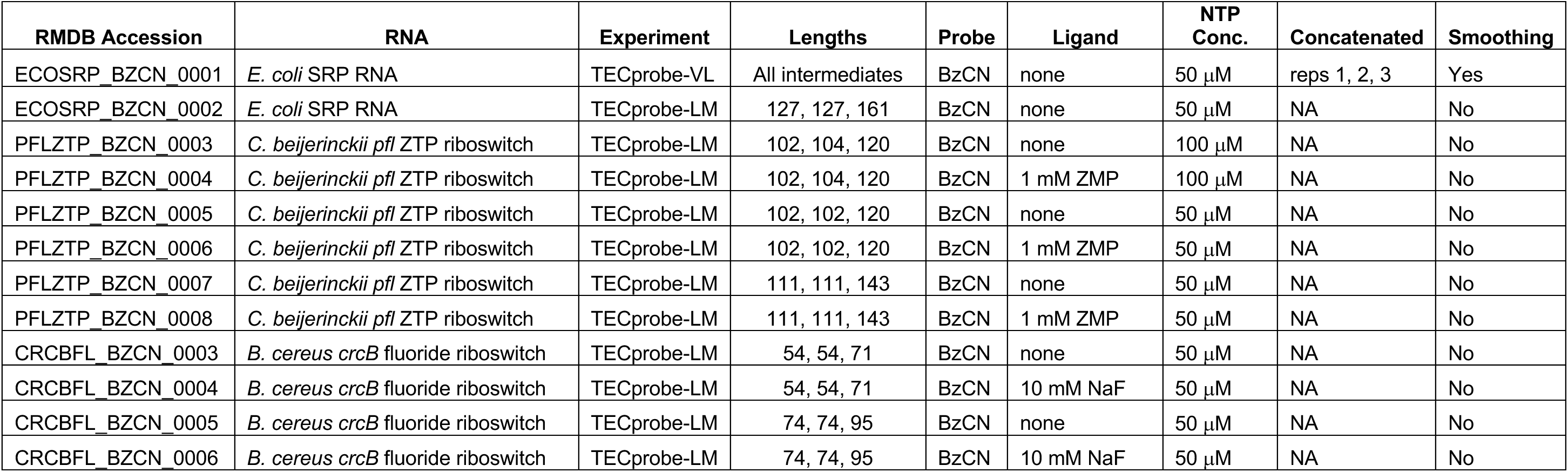
RMDB data deposition table. Reactivity data generated in this work are freely available from the RNA Mapping Database (RMDB) (http://rmdb.stanford.edu), accessible using the RMDB ID numbers indicated in the table below.

